# Distinct genomic contexts predict gene presence-absence variation in different pathotypes of a fungal plant pathogen

**DOI:** 10.1101/2023.02.17.529015

**Authors:** Pierre M. Joubert, Ksenia V. Krasileva

## Abstract

**Background:** Fungi use the accessory segments of their pan-genomes to adapt to their environments. While gene presence-absence variation (PAV) contributes to shaping these accessory gene reservoirs, whether these events happen in specific genomic contexts remains unclear. Additionally, since pan-genome studies often group together all members of the same species, it is uncertain whether genomic or epigenomic features shaping pan-genome evolution are consistent across populations within the same species. Fungal plant pathogens are useful models for answering these questions because members of the same species often infect distinct hosts, and they frequently rely on gene PAV to adapt to these hosts.

**Results:** We analyzed gene PAV in the rice and wheat blast fungus, *Magnaporthe oryzae*, and found that PAV of disease-causing effectors, antibiotic production, and non-self-recognition genes may drive the adaptation of the fungus to its environment. We then analyzed genomic and epigenomic features and data from available datasets for patterns that might help explain these PAV events. We observed that proximity to transposable elements (TEs), gene GC content, gene length, expression level in the host, and histone H3K27me3 marks were different between PAV genes and conserved genes, among other features. We used these features to construct a random forest classifier that was able to predict whether a gene is likely to experience PAV with high precision (86.06%) and recall (92.88%) in rice-infecting *M. oryzae*. Finally, we found that PAV in wheat- and rice-infecting pathotypes of *M. oryzae* differed in their number and their genomic context.

**Conclusions:** Our results suggest that genomic and epigenomic features of gene PAV can be used to better understand and even predict fungal pan-genome evolution. We also show that substantial intra-species variation can exist in these features.

## Background

Microbial species have expansive pan-genomes that allow them to adapt to their environments. While bacteria typically gain and lose genes in the form of large horizontal gene transfer events [1], the accessory portion of fungal pan-genomes, which is defined in contrast to the conserved set of genes found in all members of a species, are typically shaped by small gene duplication and deletion events, which contribute to gene presence-absence variation (PAV) [2]. Previous fungal pan-genome studies have focused on the roles and functions of core and accessory genes [2–5], but our knowledge of which genomic and epigenomic features shape fungal pangenomes remains limited. Some studies have highlighted an association of accessory genes with subterminal chromosomal regions and transposable elements (TEs) [2,3], but it is uncertain whether these associations are strong enough to be predictive of gene PAV. While one study constructed models that could successfully predict meiotically derived structural variation generation events, and identified TEs, histone marks and GC content as particularly important predictors, it did not expand these findings to pan-genome evolution [6]. Finally, it is unclear whether any patterns in genomic or epigenomic features of PAV events would be generalizable to all populations of the same species, as pan-genomes are typically assembled for entire species without consideration of differential evolution between populations.

Fungal plant pathogens are ideal candidates to study pan-genome evolution, and especially gene PAV. They have dynamic pan-genomes that allow them to adapt to their hosts [3–5]. Specifically, fungal plant pathogens secrete a wide range of rapidly evolving effector proteins to cause disease. These effectors can become a disadvantage, however, when the immune receptors of their hosts acquire new binding specificities that detect these effectors and trigger an immune response [7]. Gene PAV is therefore particularly important in fungal plant pathogen evolution [8]. In these fungi, rapid genome evolution, especially of effectors, tends to occur in TE-dense and gene-poor regions of the genome while slower evolution and house-keeping genes occur in TE-poor and gene-dense regions of the genome [9,10]. This idea is often referred to as the “two-speed” genome concept. While effectors are particularly prone to PAV, it is unclear whether this concept extends to gene PAV and especially PAV of non-effectors. Many fungal plant pathogens of the same species also infect distinct hosts, which could facilitate the characterization and comparison of these PAV events in isolated populations.

*Magnaporthe oryzae* causes the blast disease of rice and wheat and is amongst the most important and well-studied pathogens with hundreds of available genomes and next-generation sequencing datasets [11,12]. The fungus has been reported to experience substantial gene PAV but these analyses have been largely restricted to effectors, and the genomic and epigenomic features associated with these PAV events remain largely unexplored [13–15]. *M. oryzae* reproduces mostly clonally, which makes the study of how its pan-genome can evolve without substantial recombination possible [15–17]. Finally, the blast fungus infects several different hosts, enabling the comparison of gene PAV between pathotypes within the same species [17]. While rice blast has been a long-standing threat, the rapid spread of wheat blast throughout the world as well as its devastating effect on wheat crops has strongly encouraged research into this pathotype of *M. oryzae* and especially how it was able to make the jump hosts from rice to wheat and become such a devastating pathogen [12]. Altogether, *M. oryzae* offers a unique opportunity for studying gene PAV and the genomic and epigenomic features that shape these events as well as how these events vary within a species.

In this study, we sought to characterize and compare gene PAV in rice-infecting (MoO) and wheat-infecting *M. oryzae* (MoT). We first identified orthogroups experiencing PAV that distinguished isolated MoO lineages and found that they were enriched in effectors, antibiotic production, and non-self-recognition. Next, we characterized the genomic contexts in which all gene PAV occurs in MoO and MoT and found that TEs were often found in proximity to these genes. Additionally, we found that gene length, GC content, expression, and histone H3k27me3 marks were distinct for PAV genes. We used these features to construct a random forest classifier and found that the differences we observed were strong enough to produce a model that predicted whether a gene is likely to experience PAV with high precision (86.06%) and recall (92.88%). Finally, we found significant differences in the number of PAV events and the features that predict PAV in MoO and MoT, which could reflect their differing evolutionary history and could be evidence of distinct mechanisms contributing to gene loss in the two recently diverged lineages.

## Results

### Genes associated with pathogenicity, non-self-recognition and antibiotic production, are enriched among orthogroups experiencing lineage-differentiating presence-absence variation in *M. oryzae*

Differences in gene PAV events between isolated lineages of *M. oryzae* could be evidence of local adaptation. MoO isolates can be grouped into four lineages, called lineages 1, 2, 3, and 4 [17]. Lineages 2, 3, and 4 are monophyletic within the MoO phylogeny and propagate clonally [17]. All lineages show evidence of local adaptation [15]. To generate a table of all gene PAV events in MoO, we analyzed 123 previously published genomes [15]. These genomes were re-annotated, and the proteomes were clustered into orthogroups. This enabled us to identify putative gene absences in all genomes. These were then validated by using TBLASTN [18] against the genome and comparing hits to the missing orthogroup using BLASTP[18]. This approach helped ensure that gene absences were not annotation errors. We also constructed a phylogeny from a multiple sequence alignment of all of our single copy orthologs (SCOs) and found each of the three clonal MoO lineages formed separate monophyletic groups in our data, as previously observed (Additional File 1: Fig. S1) [15,17].

To identify whether differences in gene PAV events existed between the three clonal lineages of MoO, we performed a principal components analysis (PCA) on our table of PAV events. We found that the top 2 principal components (PCs) of our PCA clearly separated the lineages demonstrating that different PAV events had occurred in each lineage since their separation (Fig. 1A). Next, we identified 587 orthogroups that represented 70.53% and 62.17% of the variance in PCs1 and 2, respectively, and labeled these orthogroups as experiencing lineage-differentiating PAV. We then identified, among all orthogroups, 594 putative effector orthogroups and found, as previously reported [14,15], that PAV of effector orthogroups alone were sufficient to separate the MoO lineages in a follow-up PCA (Fig. 1B). Given that we identified 4.30% of all orthogroups as putative effectors, the fact that 8.67% of lineage-differentiating PAV orthogroups were effectors represented a clear enrichment. However, non-effector orthogroups still represented 91.33% of lineage-differentiating PAV orthogroups, showing that many orthogroups besides effectors experience lineage-differentiating PAV (Fig. 1C).

**Fig. 1.**
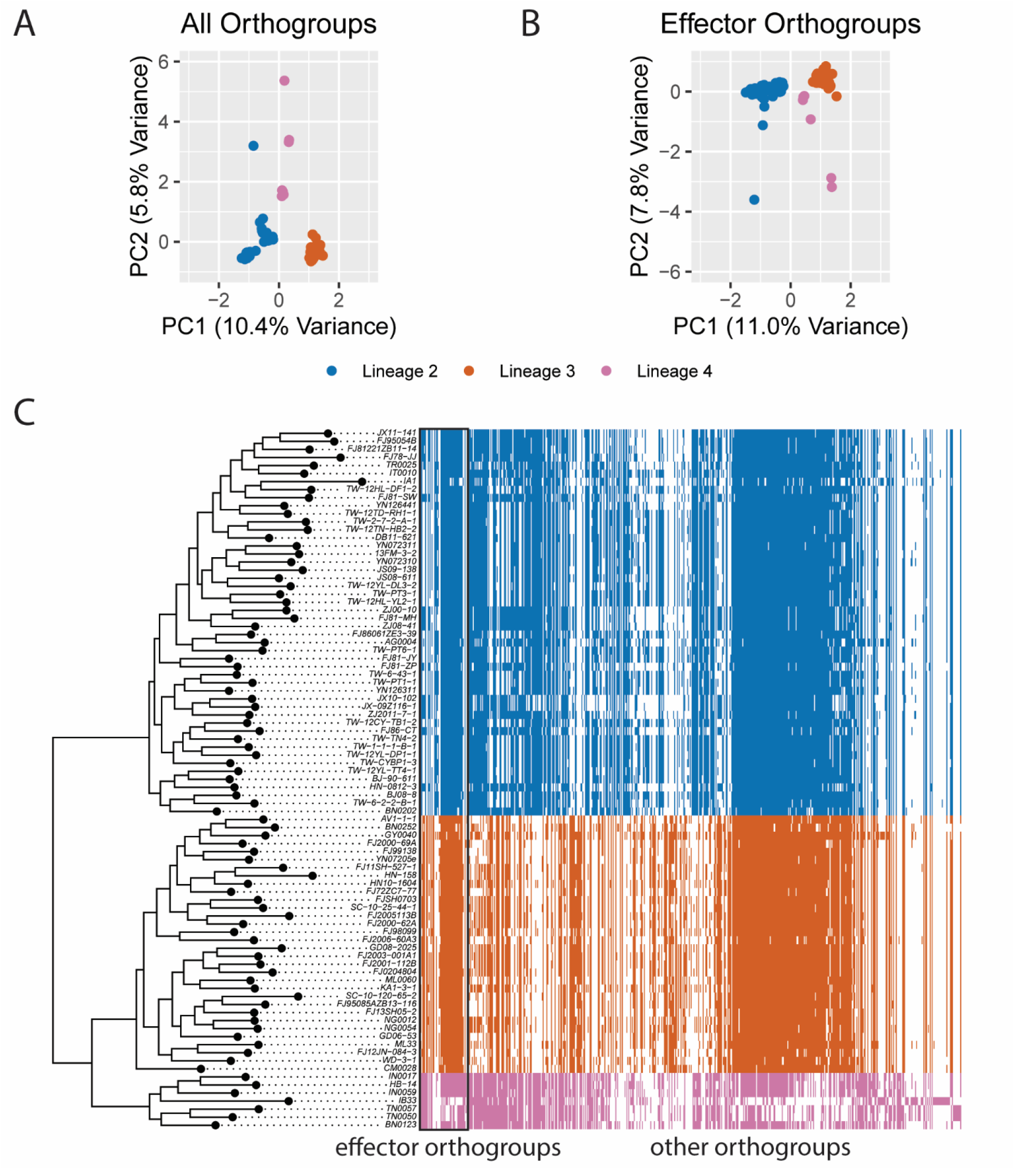
PAV of effector and non-effector orthogroups differentiate the clonal lineages of rice-infecting *M. oryzae*. A. Scatter plot of values for principal components (PCs) 1 and 2 resulting from a PCA of orthogroup PAV. Each point represents one isolate. B. Scatter plot of values for PCs 1 and 2 resulting from a PCA of effector orthogroup PAV. Each point represents one isolate. C. Heat map representing which orthogroups are present (color) or absent (white) in each genome. Effector orthogroups are separated from other orthogroups by a black box. The phylogeny was generated using a multiple-sequence alignment of SCOs and fasttree and is a subset of the full MoO phylogeny generated from our data (Additional File 1: Fig. S1). In all panels, colors represent the clonal lineages of MoO. Blue represents lineage 2, orange represents lineage 3 and pink represents lineage 4. Lineages were named as previously described [17].

To identify what other types of genes were enriched amongst lineage-differentiating PAV orthogroups, we performed gene ontology (GO) and protein family (PFAM) enrichment analysis. This analysis revealed that lineage-differentiating PAV orthogroups were enriched for GO terms related to secondary metabolite production and biosynthesis of membrane components, among other terms (Fig. 2A). Lineage-differentiating PAV orthogroups were enriched for PFAM domains related to antibiotic production, among other domains (Fig. 2B). Genes without PFAM domains were also strongly enriched in PAV orthogroups (6040 annotated, 407 observed, 256.55 expected, p-value < 0.001, Fisher’s exact test). Notably, the HET domain, which is associated with heterokaryon incompatibility in fungi, was also enriched among these orthogroups (Fig. 2B). Finally, while NACHT and NB-ARC domains did not appear enriched on their own due to a small number of lineage-differentiating PAV orthogroups having these annotations, NOD-like receptors (NLRs) which may play a import role in fungal immunity and contain either a NACHT or an NB-ARC domain [19], were enriched amongst lineage-differentiating PAV orthogroups (23 annotated, 4 observed, 1.15 expected, p-value = 0.026, Fisher’s exact test). These results indicated that antibiotic production and non-host recognition, in addition to effectors, may play an important role in driving adaptation in these three isolated lineages of rice-infecting *M. oryzae*.

**Fig. 2.**
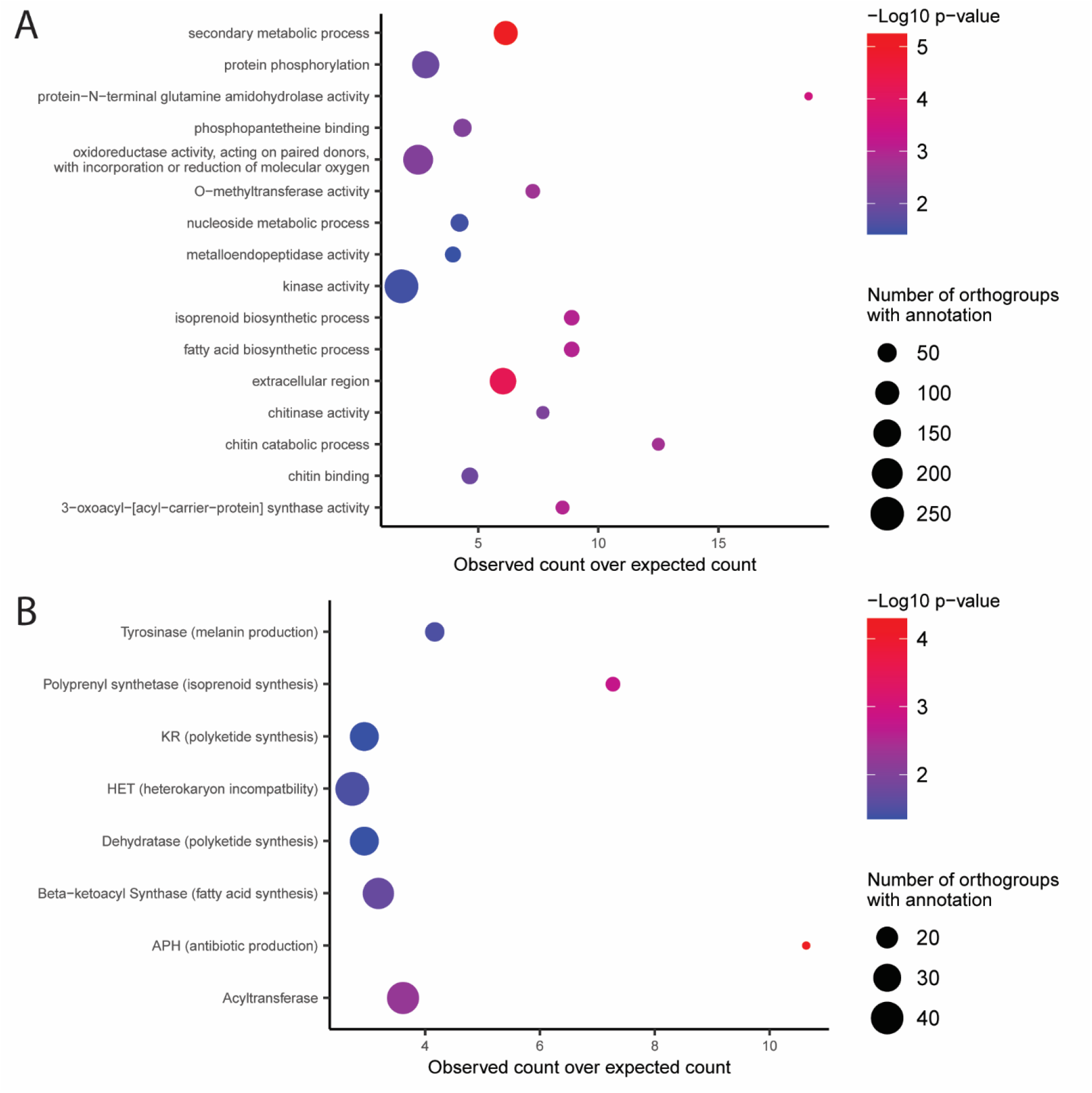
Lineage-differentiating PAV orthogroups in rice-infecting *M. oryzae* contain many genes related to antibiotic production and non-self-recognition. A. Gene ontology (GO) enrichment analysis of lineage-differentiating PAV orthogroups. B. Protein family (PFAM) domain enrichment analysis of lineage-differentiating PAV orthogroups. P-values shown are the results of Fisher’s exact tests.

### Presence-absence variation genes are more common and more spread out throughout the genome in wheat-infecting *M. oryzae* than in rice-infecting *M. oryzae*

We next sought to identify whether there were specific patterns in the genomic contexts of PAV events in *M. oryzae*. To expand our analyses beyond lineage-differentiating PAV orthogroups and to compare PAV orthogroups to conserved orthogroups, we first needed a systematic way to label them. To avoid erroneously calling single gene gain or loss events as PAV, we chose to incorporate phylogenetic information in these definitions and therefore identified PAV and conserved orthogroups for each clonal, monophyletic lineage of *M. oryzae*. In our data, orthogroups were labeled as PAV if they were present in all isolates of at least two subclades within a lineage and absent in all isolates of at least two subclades within a lineage. This definition meant that at least two phylogenetically independent loss or gain events needed to be observed in our data for an orthogroup to be labeled PAV. All orthogroups that were present in all but two or fewer isolates in a lineage were labeled as conserved orthogroups. All orthogroups that did not fit this definition were labeled as “other”. Genes belonging to PAV orthogroups or conserved orthogroups were labeled PAV genes and conserved genes, respectively. This approach allowed us to label 1,269 and 1,029 PAV orthogroups in lineage 2 and 3 of MoO, respectively (Fig. 3A). We did not include lineage 4 in our analysis because of its small sample size and omitted lineage 1 because it is thought to be recombining and would therefore would have violated the assumptions of our definition of PAV and conserved orthogroups [17].

**Fig. 3.**
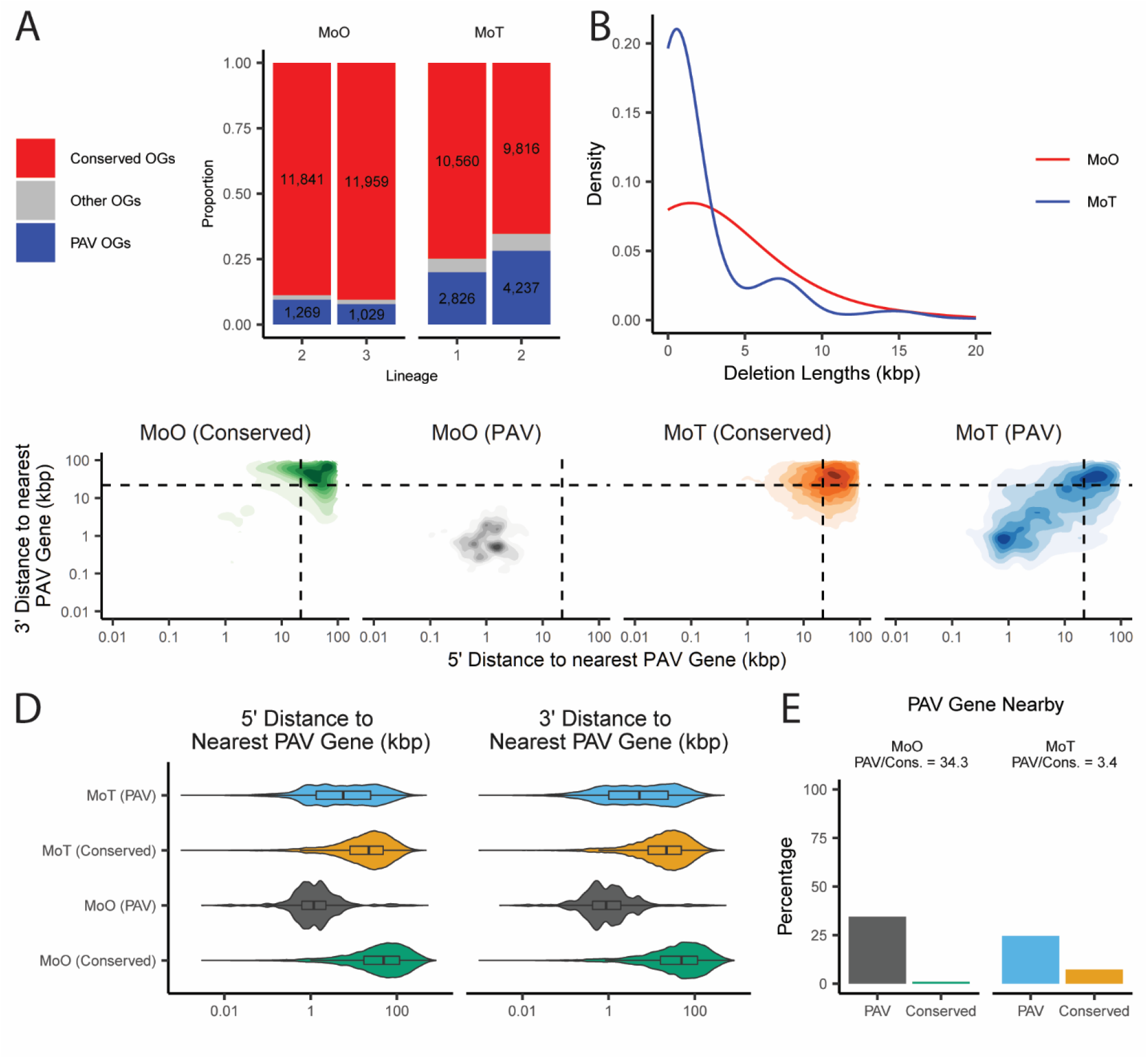
PAV genes are more common and more spread out throughout the genome in wheat-infecting *M. oryzae* than in rice-infecting *M. oryzae*. A. Stacked barplot comparing the number of PAV orthogroups (OGs) and conserved orthogroups in MoO and MoT. “Other OGs” denote orthogroups that did not satisfy our definitions for either category. B. Distribution of the lengths of genomic deletions in MoO and MoT. C. Density plot showing the distribution of the distances to the nearest PAV gene for conserved and PAV genes in MoO and MoT. Dashed lines in density plots represent the median values for all genes in both pathotypes. D. Violin plot showing the distribution of the distances to the nearest PAV gene for conserved and PAV genes in MoO and MoT. E. Percentages and proportions of PAV and conserved genes that are within 1000bp of a PAV gene in in MoO and MoT. Median values and statistical comparisons for data shown in panels C through E are listed in Additional File 7, Additional File 8, and Additional File 9.

To compare PAV across rice and wheat-infecting *M. oryzae* isolates, we annotated, called orthogroups and validated missing orthologs for 88 previously published MoT genomes (Additional File 2). Unlike for MoO, MoT isolates have not been formally assigned into lineages in the past, though they are also thought to have propagated mostly clonally since their recent appearance [16]. We therefore generated a phylogeny using our SCOs and identified two monophyletic MoT lineages in our data, which we called lineages 1 and 2 (Additional File 1: Fig. S2). MoT lineages 1 and 2 were made up of 39 and 48 isolates, respectively, meaning these lineages were similar in size to the previously described MoO lineages (Additional File 1: Fig. S1). In the two MoT lineages, we identified more than twice the number of PAV orthogroups than in the MoO lineages (Fig. 3A). To assess whether this contrast was also present for genomic deletions, we used 117 MoO and 47 MoT Illumina whole-genome sequencing datasets to call genomic deletions based on a high-quality reference genome for each pathotype (Additional File 3 and Additional File 4). This approach allowed us to identify 1,870 deletions in MoO and 1,862 deletions in MoT despite using more than double the number of datasets for MoO than MoT (Additional File 5 and Additional File 6). We also found that genomic deletions were larger in MoO than in MoT, with a median of 1,818bp in MoO and 960bp in MoT (Fig. 3B). Correspondingly, when we compared the density of PAV genes in MoO and MoT, we found that genes belonging to PAV orthogroups were much more likely to be in proximity with other genes belonging to PAV orthogroups in MoO than in MoT (Fig. 3C-E, Additional File 7, Additional File 8, and Additional File 9). Taken together, these results indicated that genomic deletions, especially those involving genes, were more likely to involve multiple genes in MoO than in MoT. These results also hinted that gene PAV happens in defined regions of the genome in MoO while in MoT these events are more likely to be randomly spread out throughout the genome.

### Genes prone to presence-absence variation are closer to transposable elements than other genes

The two-speed genome hypothesis defines two genomic compartments in fungal plant pathogens, one characterized by rapid evolution, few genes and many TEs, and the other characterized by slow evolution, many genes and few TEs [9,10]. We investigated whether orthogroups experiencing PAV followed this model in *M. oryzae*. We found that genes in PAV orthogroups were much closer to TEs than genes in conserved orthogroups in both MoO and MoT (Fig. 4, Additional File 7 and Additional File 8). While the differences in distance to the nearest gene between conserved and PAV orthogroups in MoO or MoT were typically quite small (median difference <100bp), we did find that genes in PAV orthogroups were less likely to be close to genes than conserved genes, though the effect was not as strong as for TEs. (Additional File 1: Fig. S3, Additional File 7, and Additional File 8). We also observed differences in these patterns for MoO and MoT. Specifically, we found that PAV orthogroups in MoO were more likely to be close to TEs than those in MoT (Fig. 4C and Additional File 9). We also found that MoO PAV genes were more likely to be far away from genes that MoT PAV genes (Additional File 1: Fig. S3C and Additional File 9).

**Fig. 4.**
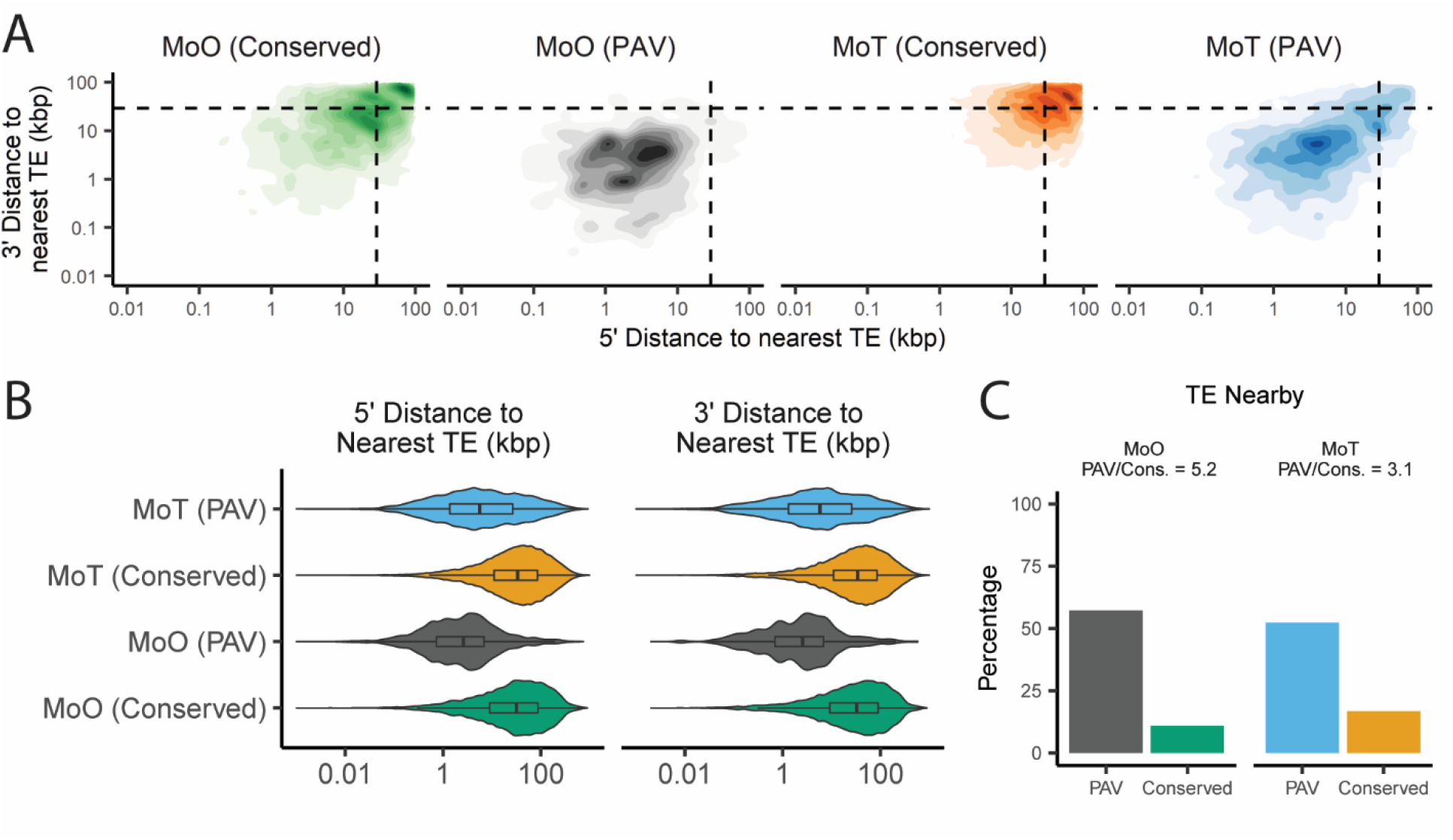
PAV genes are more likely to be found near transposable elements (TEs) than conserved genes. A. Density plots showing the distribution of the distances to the nearest TE for conserved and PAV genes in MoO and MoT. B. Violin plot showing the distribution of the distances to the nearest TE for conserved and PAV genes in MoO and MoT. C. Percentages and proportions of PAV and conserved genes that are within 5000bp of a TE in MoO and MoT. Dashed lines in density plots represent the median values for all genes in both pathotypes. Median values and statistical comparisons for data shown are listed in Additional File 7, Additional File 8, and Additional File 9.

To understand if these observations also applied to genomic deletions in MoT and MoO, we measured TE and gene densities within the genomic deletions we previously identified and within their flanking regions. This analysis revealed that genomic deletions and their flanking regions were enriched in TEs and depleted in genes, though the effect was stronger for TE density than for gene density (Additional File 1: Fig. S4).

### Genes prone to presence-absence variation show distinct genomic and epigenomic features and some differences in these features between rice and wheat-infecting *M. oryzae*

A previous report has shown that MoO has a much greater TE content than MoT [20]. Therefore, given this result and the greatly increased number of PAV orthogroups in MoT compared to MoO we observed (Fig. 3A), it is unlikely that TEs alone define whether a gene is prone to PAV or not. We next chose to investigate whether we could identify other differences in genomic features between PAV genes and conserved genes in *M. oryzae*. We first looked at the GC content of these genes and the regions that flank them. PAV genes were more likely to have lower GC content than conserved genes (Fig. 5A and Additional File 10), as did the regions that flank them, though the effect was more subtle for the flanking regions (Additional File 1: Fig. S5A and Additional File 10). We also found that PAV genes were shorter than conserved genes (Fig. 5B and Additional File 10). We next performed various functional annotations of PAV and conserved genes and found that PAV genes were more likely to be predicted effectors, in accordance with our previous results, and less likely to have GO or PFAM annotations than conserved genes (Additional File 1: Fig. S6 and Additional File 8).

**Fig. 5.**
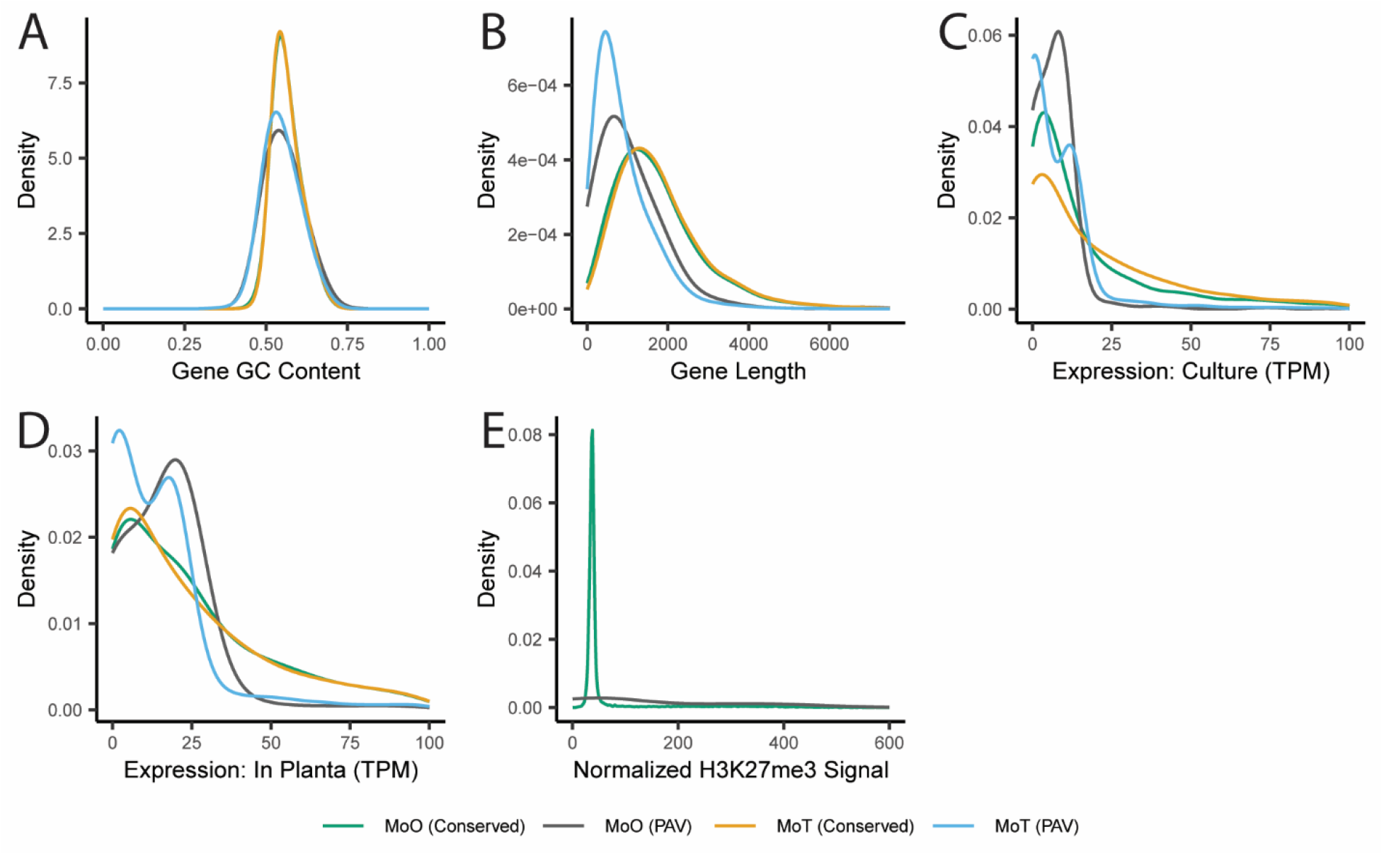
PAV genes are distinct from conserved genes in many ways beyond their proximity to TEs. Density plots showing the distributions of A. gene GC content, B. gene lengths, C. expression in culture, D. expression *in planta*, and E. normalized H3K27me3 histone mark ChIP-Seq signal for PAV and conserved genes in MoO and MoT. In panel A, the line representing the data for conserved genes in MoO appears underneath the line representing the data for conserved genes in MoT. In panel E, MoT genes were not included as this data is not available for MoT. Statistics describing the distributions shown and statistical comparisons between these statistics are listed in Additional File 8, Additional File 9, Additional File 10, and Additional File 11.

Next, we gathered histone mark, transcription, methylation, and extrachromosomal circular DNA sequencing (eccDNA) data from the literature for both MoO and MoT to further characterize PAV genes. Unfortunately, these datasets were only available for some strains of MoO or MoT but not for all. Therefore, we analyzed these datasets for one reference MoO strain and one reference MoT strain, and then generalized this signal to our orthogroups to impute the signal in other strains of *M. oryzae*. This allowed us to observe that average expression was higher both in culture and *in planta* for conserved genes than for PAV genes (Fig. 5C and D and Additional File 10). Additionally, PAV genes were more likely to show signal from chromatin immunoprecipitation sequencing of H3K27me3 and H3K36me3 histone marks and less likely to show signal from H3K27ac histone marks (Fig. 5E, Additional File 1: Fig. S5B and C, and Additional File 10). We also looked at bisulfite sequencing data and found that PAV genes were less methylated and showed a greater variation in methylation percentage than conserved genes (Additional File 1: Fig. S5D and Additional File 10). Finally, we found that PAV genes had a much tighter distribution of eccDNA sequencing signal than conserved genes (Additional File 1: Fig. S5E and Additional File 10). Overall, these results indicated clear differences in the genomic and epigenomic features of PAV genes compared to conserved genes.

Many differences between PAV genes in MoO and MoT were discovered through this process including in gene length, where PAV genes were smaller in MoT than MoO (Fig. 5B and Additional File 11), and in expression, where PAV genes in MoT showed a bimodal distribution and less expression on average than MoO PAV genes both in culture and *in planta* (Fig. 5C and D and Additional File 11). Additionally, PAV genes were more likely to have GO and PFAM annotations in MoO than in MoT (Additional File 1: Fig. S6E and F and Additional File 9). These observations further supported the idea that PAV may be occurring in different genomic contexts in MoO and MoT.

Finally, we analyzed a similar set of features in the genomic deletions we identified in MoT and MoO. We found that while some of the trends we observed in PAV genes were similar in genomic deletions compared to a genomic baseline, such as H3K27me3 signal and GC content, other trends like the increase in expression of PAV genes did not translate to differences in RNAseq signal in deleted regions (Additional File 1: Fig. S7 and Additional File 12).

### Genomic and epigenomic features of genes prone to presence-absence variation can be used to generate predictive models for rice and wheat-infecting *M. oryzae*

Our previous results demonstrated the differences in genomic contexts between PAV genes and conserved genes. We then wanted to determine whether these features in aggregate could provide enough signal to predict whether a gene was prone to PAV using a machine learning approach. To this end, we trained a random forest classifier on all features we described for MoO. We selected this algorithm because of its ease of implementation as well as its robustness to correlated features [21]. When we trained this model on data from all but 8 strains of MoO and tested the model on the remaining strains, we observed that the model performed very well and was able to predict PAV genes with 86.06% precision and 92.88% recall on average (F1 = 89.34%, Fig. 6A). Our model also allowed us to determine how important each feature was in predicting PAV genes by calculating the decrease in the F1 statistic when the variable in our testing data was permuted. This approach identified histone H3K27me3 as being the most predictive feature for PAV genes in MoO (Fig. 6B). Although the accuracy of predictions by the random forest classifier is robust to correlated features, the variable importances we observed were likely influenced by the fact that several variables in our model were correlated with each other and that many showed high dependences, which meant that the information encoded in these variables could also be described by other variables in the model (Additional File 1: Fig. S8 and Fig. S9). Next, we trained a model to predict PAV genes in MoT and found that the model performed even better with a precision of 94.25% and a recall of 96.67% (F1 = 95.44%, Additional File 1: Fig. S10A). In this reduced model, gene expression in culture stood out as being particularly predictive of MoT PAV genes (Fig. 6C). Finally, we trained another MoO model using a reduced set of features that matched the data that was available for MoT, and found that the MoO model still performed well with an 86.11% precision and 92.21% recall (F1 = 89.05%, Additional File 1: Fig. S10B). The similar performance of the two MoO models could likely be explained by the high dependences of our variables (Additional File 1: Fig. S8 and Fig. S9). When comparing the reduced MoO model to the MoT model, we noticed some differences between the importances of the features in each model (Fig. 6C and D). For example, in culture expression and the presence of functional annotations being more important in the reduced MoO model than in the MoT model. These differences in importances may have been influenced by the previously described differences in the features of PAV genes in MoO and MoT.

**Fig. 6.**
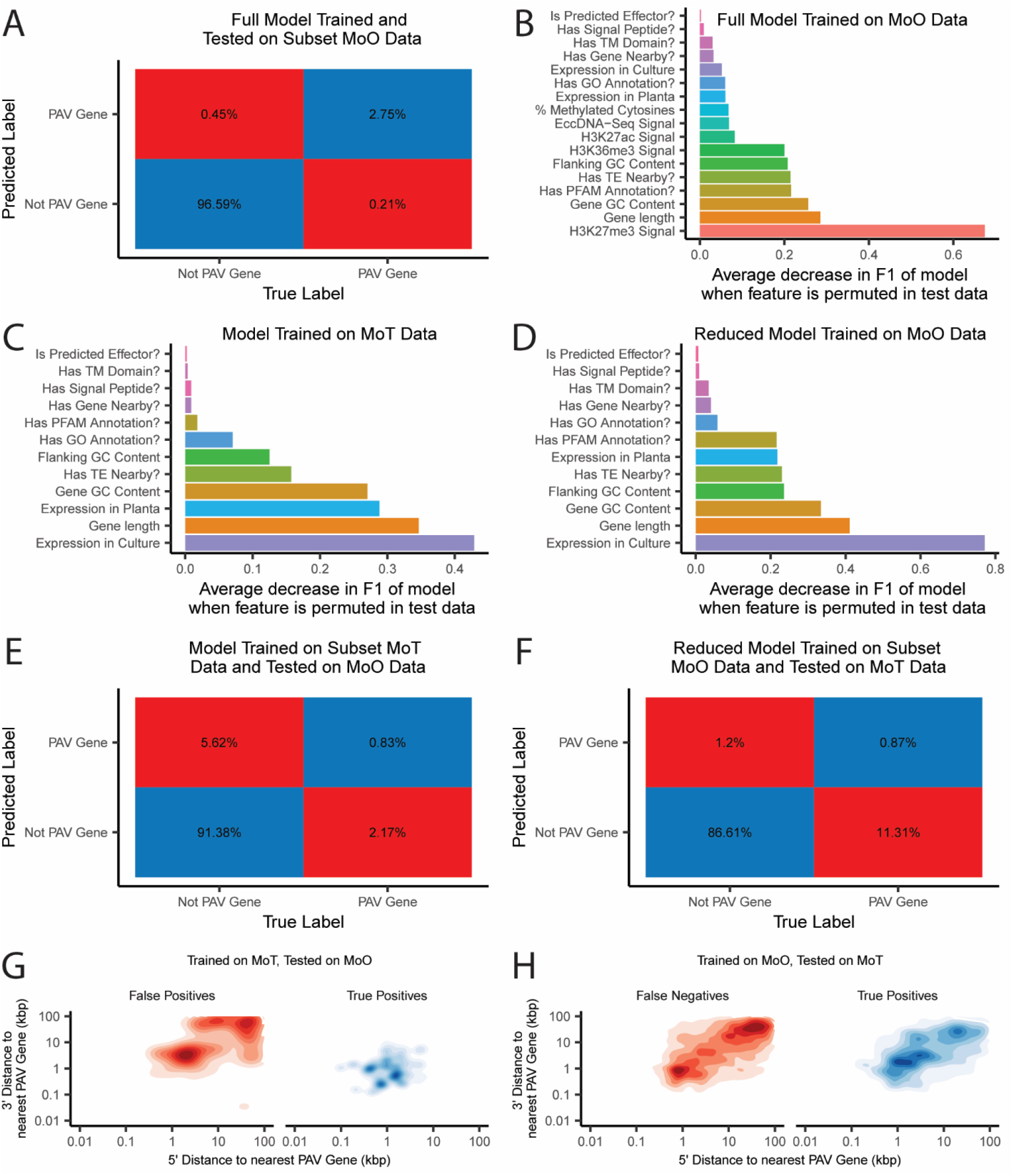
Random forest classifiers accurately identify PAV genes in rice and wheat-infecting *M. oryzae*, but the models perform poorly on genes from the host they were not trained on. A. Confusion matrix showing predictions of the MoO random forest classifier when tested on MoO genes that it was not trained on. B. Decrease in the F1 statistic of the MoO random forest classifier when each feature is permuted in the testing data. Features described as questions are binary, all other features are continuous. C. Decrease in the F1 statistic of the MoT random forest classifier when each feature is permuted in the testing data. D. Decrease in the F1 statistic of the MoO random forest classifier trained on a subset of features when each variable is permuted in the testing data. F. Confusion matrix showing predictions of the MoT random forest classifier when tested on MoO genes. E. Confusion matrix showing predictions of the MoO random forest classifier trained on a reduced subset of features when tested on MoT genes. G. Density plots showing the distribution of the distances to the nearest PAV gene for false positive and true positive predictions by the MoT random forest classifier when tested on MoO genes. H. Density plots showing the distribution of the distances to the nearest PAV gene for false negative and true positive predictions by the MoO random forest classifier trained on a subset of features when tested on MoT genes.

### A predictive model trained on wheat-infecting *M. oryzae* data does not accurately predict presence-absence variation in rice-infecting *M. oryzae* and vice versa, highlighting differences in the genomic contexts of presence-absence variation in the two pathotypes

Finally, we tested if the model trained on MoT data could predict whether genes are prone to PAV in MoO and vice versa. To our surprise, the MoT model performed very poorly on MoO data, with a precision of 12.86% and a recall of 27.65% (F1 = 17.55%,Fig. 6E). Similarly, the reduced MoO model performed very poorly on the MoT data with a precision of 42.04% and a recall of 7.15% (F1 = 9.55%, Fig. 6E). Differences in the importances of the features in each model likely provided some explanation for this observation (Fig. 6C and D). Additionally, we noticed that the MoO model had predicted too few PAV genes in the MoT data and that the MoT model had predicted too many PAV genes in the MoO data (Fig. 6E and F). When we analyzed the conserved genes that the MoT model falsely labeled as PAV, we found that many of them were found in isolated regions far away from true PAV genes (Fig. 6G). Similarly, many of the PAV genes in MoT that were not detected by the MoO model were found in isolated regions (Fig. 6H). These results followed the patterns we observed in PAV clusters in MoO and MoT genomes (Fig. 3A). Our observations, combined with the differences we observed in the genomic and epigenomic features of PAV genes in MoO and MoT described previously, indicated that the patterns and genomic contexts of PAV between the two pathotypes are significantly different, despite being within the same species.

## Discussion

Gene PAV plays an important role in fungal pan-genome evolution [2–5]. To improve our understanding of these events, we designed a robust pipeline to identify orthogroups experiencing PAV in *M. oryzae*. We found that PAV of these orthogroups differentiates isolated lineages of MoO, and found that these lineage-differentiating PAV orthogroups are enriched for effectors, as previously published [14,15]. We also found that genes related to antibiotic production and non-self-recognition were also enriched among them. This result could point to the local and rice-associated microbiome playing an important role in *M. oryzae*’s evolution. All three clonal lineages are geographically isolated and experience different climates [15]. They also tend to infect different rice varieties and cause disease of varying severity [15]. Geography and host genotype could have major influences on the microbiome the fungus encounters. Microbiome sampling of rice varieties used in these areas as well as the environment could therefore give better insight into the results we present here and how the microbiome might shape the fungus’s fitness. Additionally, it is important to note that adaptation to the host microbiome and the environment in general are often forgotten when discussing fungal plant pathogen evolution. Our results point to the importance of considering these factors when studying the success of these pathogens. Unfortunately, we could not extend these analyses to MoT as lineages of MoT have not been formally defined in the past and neither has their geography or host phenotypes.

We then looked to find features of PAV orthogroups that might help us better understand where these events are occurring in the genome. We found that these events were associated with a high TE density and a low gene density, though the effect was stronger for TE density than gene density. We also found that PAV genes are shorter, have lower GC content, and are more likely to be effectors. Finally, PAV genes are more highly expressed and display stronger histone H3K27me3 signal than conserved genes. When we combined all of these features into a predictive model, we found that the model performed very well and predicted PAV genes with 86.06% precision and 92.88% recall, on average. We were also able to identify histone H3K27me3 as the most predictive feature, though gene length and GC content stood out as well. We could not clearly state whether genomic deletions showed similar features to PAV genes in our data. While we found that these genomic deletions occurred frequently in TE-dense and gene-sparse areas of the genome, and that GC content and H3K27me3 ChIP-Seq signal for these regions resembled that of PAV genes, other features were not similar between the two. These results may have been confounded by the need for reference-based identification of these deletions, an unclear baseline for comparison, and events like transposon insertion polymorphisms.

Many of the features that were particularly important in our classifier were related to the two-speed genome concept which supported the idea that gene PAV in *M. oryzae* is strongly associated with the rapidly evolving compartment of the genome [9,10]. Our findings support the idea that these features may play an important role in the evolution of the pathogen. However, the fact that the presence of TEs were important features in our random forest classifier but not amongst the most important, supports the idea that the correlation between TEs and rapid evolution is not always a causal one and that complex correlations are at play. In short, other variables may be shaping the PAV-prone compartment of the *M. oryzae* genome and driving both rapid evolution and TE activity. Our findings also reflect previous findings on the association between TEs and the evolution of the accessory portion of fungal pan-genomes [3]. While the gene space for the genomes we analyzed were well assembled, most of the genomes we performed our analysis on were not chromosome-level assemblies. Therefore, although we observed features associated with subterminal regions in our PAV genes, we could not confirm previous findings on the association between subterminal regions and accessory genes in other fungi [2]. This analysis should be repeated once more high-quality genomes become available for *M. oryzae* to fully determine whether these findings apply to the blast fungus as well. Similarly, though the features we identified in this study should be kept in mind when studying other pan-genomes, it is unclear whether the features of gene PAV we identified are applicable to other fungi, and therefore more in-depth studies of these genomic and epigenomic features are necessary to assess how broad these findings are as datasets become available for more fungi.

Though our random forest classifier performed well, many of the important features we used in our model had to be propagated from a single strain which likely led to many biologically inaccurate values in our data and potentially errors in how we ranked the importances of each variable. To truly validate our results, our approach would need to be repeated using more data for each isolate. However, even a model using a subset of our features performed well, indicating that RNAseq data for each isolate alone may be enough to further substantiate these results. Regardless, our model showed that PAV genes can be identified simply using features in the genome, therefore establishing a method to identify genes prone to PAV in *M. oryzae* without relying on phylogenetics. This could be useful for identifying genes prone to PAV in lineages of MoO with very few isolates, like lineage 4, or for studying PAV in groups of genes with complicated evolutionary relationships like sequence unrelated structurally similar (SUSS) effectors [22]. While our models performed well, they also identified many genes that had features of PAV genes but did not experience PAV. These false positives could help us better understand which genes are under strong selection to be kept in the *M. oryzae* genome or which genes’ genomic contexts are changing to look more like conserved genes. Notably, our results support the exciting possibility of using genomics to predict targets for disease-prevention strategies that will remain in the genome, therefore making these strategies more robust.

Finally, we found distinct patterns in the genomic contexts of PAV genes in MoO and MoT. Specifically, we found that PAV in MoT appeared to occur more frequently and was more spread out throughout the genome than in MoO. Though we were able to find two lineages of MoT in our data that contained similar numbers of isolates to the previously described lineages of MoO [17], the evolutionary distances between isolates in lineages of the two pathotypes were quite different, which may have contributed to the differences in the number of PAV orthogroups we observed (Additional File 1: Fig. S1 and Fig. S2). However, the strength of these differences, as well as supporting evidence from our analysis of genomic deletions, suggest that our observations are valuable despite this caveat. We also found that many of the genomic and epigenomic features of PAV that we identified in MoO were different in MoT. These differences culminated in our MoO random forest classifier performing poorly on MoT data and vice versa, with specific patterns in the false positives and false negatives of these tests which reflected the observed differences between MoO and MoT. These results in aggregate indicated differences in the evolution of the rice and wheat pathotypes of *M. oryzae*.

The two *M. oryzae* pathotypes share some major differences in their TE content [20] and very different life histories, with MoO originating 9,800 thousand years ago [17] and propagating mostly clonally since then, while MoT is thought to have emerged approximately 60 years ago from a multi-hybrid swarm of many different *M. oryzae* pathotypes [12,16]. We propose that the differences in PAV across the two pathotypes may reflect these life histories, with MoO exhibiting more of a stable equilibrium and much slower paced evolution, where PAV events happen in specifically defined compartments of the genome, while MoT is rapidly losing and gaining genes, even in areas of the genome where most of the conserved genes in MoO are located. It is unclear at this point whether MoT is headed towards an equilibrium that will resemble MoO, or whether there are key differences between the two pathotypes that are shaping their genomes beyond their evolutionary histories. MoT, which appears to lose genes at a faster rate than MoO and evolve rapidly in general, will pose a significant challenge for disease prevention. A better understanding of these evolutionary dynamics and the differences between MoO and MoT could help us better comprehend why MoT is such a devastating emerging pathogen and help us curb its threat. Finally, these results highlight the need to study isolated populations of a species separately as well as in aggregate to understand whether observations made for the pan-genome applies to every population within a species, especially if they are adapted to different hosts or environments and if they have different evolutionary histories.

## Conclusions

Our study demonstrates that gene PAV can be associated with specific genomic and epigenomic features in fungi and that these associations can be predictive. We also show that major variation can exist in these features between different populations of the same species. This study therefore highlights the need for more studies of fungal pan-genomes and the genomic and epigenomic features that define them to better understand how fungi adapt to their environments. These studies could also lead to a greater understanding of how fungal plant pathogens adapt to their hosts. Predicting these adaptations could help us develop more effective disease prevention strategies in the future. Finally, it is important that future pan-genome studies be done in a way that considers intra-species variation and evolutionary history of different populations to avoid generalizing based on a reference strain or pathotype.

## Methods

### Genome annotation, proteome orthogrouping, and phylogeny

The set of 123 MoO genomes were obtained from a previously published study [15], while 88 MoT genomes (Additional File 2) as well as a single *M. grisea* proteome (GCA004355905.1) were obtained from NCBI’s GenBank. All genomes were verified to have more than 90% completeness using BUSCO version 5.2.2 and the “sordariomycetes_odb10” option [23]. Genomes were annotated using FunGAP [24] version 1.1.0 and RNAseq data obtained from Sequence Read Archive (SRA) accession ERR5875670. The “sordariomycetes_odb10” option was used for the busco_dataset argument and the “magnaporthe_grisea” option was used for the augustus_species argument. For repeat masking, a TE library generated by combining the RepBase [25] fngrep version 25.10 with a *de novo* repeat library, generated by RepeatModeler [26] version 2.0.1 run on the *M. oryzae* Guy11 genome (GCA002368485.1) with the LTRStruct option, was used for all genomes. Annotated proteomes were then used as input for OrthoFinder [27] version 2.5.4 to form two separate sets of orthogroups, one for MoO genomes and one for MoT genomes. The *M. grisea* proteome was included in both as an outgroup. Orthogrouping was performed using the “diamond_ultra_sens” parameter for sequence search, the “mafft” parameter for species tree generation and the “fasttree” parameter for gene tree generation. Single copy orthologs (SCOs) were then obtained from the OrthoFinder output, aligned using mafft [28] version 7.487 with the --maxiterate 1000 parameter and the --globalpair parameter, concatenated, and then trimmed using trimal [29] version 1.4.rev22 and a 0.8 gap threshold parameter. Finally, fasttree [30] version 2.1.10 with the gamma parameter was used to generate a phylogeny and ape [31] version 5.5 was used to root each tree on the *M. grisea* outgroup.

### Gene absence validation

A preliminary set of missing orthogroups in each genome was obtained from the OrthoFinder outputs. Gene absences were validated by first using TBLASTN [18] version 2.7.1+ with the -max_intron_length 3000 parameter to align all protein sequences from an orthogroup to the genome that was missing that orthogroups. Any orthogroup that resulted in two or more hits above 55% sequence identity, 55% query coverage and an e-value smaller than 10^−10^ when aligned to the target genome were selected for further verification. These cutoffs were optimized so that less than 1% of orthogroups were misclassified as absent in a testing set of orthogroups that were known to be present in a target genome. Finally, TBLASTN hits were extracted as protein sequences using agat_sp_extract_sequences.pl version 0.9.1 from the AGAT toolkit (https://github.com/NBISweden/AGAT), and aligned against all protein sequences in all orthogroups using BLASTP [18] version 2.7.1+. The top 100 hits were collected, and majority vote was used to determine which orthogroup the TBLASTN hit would have belonged to had it been annotated by FunGAP. If no TBLASTN hits were found or if the BLASTP hits did not match the original missing orthogroup, the absence was counted as a validated absence, otherwise it was removed from the preliminary set of missing orthogroups.

### Effector annotation

Effectors were predicted in all proteomes by first selecting genes with signal peptides which were predicted using SignalP [32] version 4.1 using the “euk” organism type and using 0.34 as a D-cutoff for both noTM and TM networks. Genes with predicted transmembrane domains from TMHMM [33] version 2.0c were then excluded. Finally, EffectorP [34] version 3.0 was used to predict effectors from this secreted gene set. Effector orthogroups were then called if at least half of the orthologs within the orthogroup were annotated as predicted effectors.

### Principal component analysis and identification of lineage-differentiating PAV orthogroups

The matrix of missing effector orthogroups for each MoO isolate was used for PCA using the prcomp function in R version 3.6.1. PCA was performed a second time using the matrix of all missing orthogroups. Lineage-differentiating PAV orthogroups were then chosen by selecting the orthogroups that contributed more than 0.1% of the variance to PCs 1 and 2 using the get_pca_var function in R version 3.6.1.

### Gene ontology and protein family enrichment analyses

All proteins were annotated for GO terms using the PANNZER2 [35] webserver and command line software SANSPANZ version 3 in October 2022. Only annotations with a positive predictive value greater than 0.6 and an ARGOT rank of 1 were kept. All GO terms assigned to genes within an orthogroup were then transferred to their orthogroup. GO term enrichment analysis was then performed using TopGO [36] version 2.36.0 and enrichment was calculated using the Fisher’s exact test and the “weight” algorithm. Only GO terms that were assigned to 3 or more lineage-differentiating PAV orthogroups and who’s enrichment was significant at a p-value of less than 0.05 were reported. PFAM enrichment analysis was performed by annotating PFAM domains using pfam_scan.pl [37] version 1.6-4 and the PFAM-A database. The output from pfam_scan.pl was parsed using K-parse_Pfam_domains_v3.1.pl (https://github.com/krasileva-group/plant_rgenes) [38] and an e-value cutoff of 0.001, and domain names were simplified by removing numbers and additional letters attached to domain names. Orthogroups were called as containing a domain if at least half of their orthologs had that domain annotation. Fisher’s exact test for enrichment was performed using the scipy.stats Python module [39] version 1.9.0. Only domains which were observed in three or more lineage-differentiating PAV orthologs and with enrichment p-values less than 0.05 were reported.

### Identification of genomic deletions

Illumina sequencing data was obtained from 117 datasets for MoO and 47 datasets for MoT from the SRA (Additional File 3 and Additional File 4). Reads were mapped to the *M. oryzae* Guy11 genome (GCA002368485.1) for MoO datasets and to the *M. oryzae* B71 genome (GCA004785725.2) for MoT datasets using BWA MEM [40] version 0.7.17-r1188. Read duplicates were marked using Picard (https://broadinstitute.github.io/picard/) version 2.9.0. Structural variants were then called using smoove (https://github.com/brentp/smoove) version 0.2.8, wham [41] version 1.7.0-311-g4e8c, Delly [42] version 0.9.1, and Manta [43] version 1.6.0 using default settings. The Delly output was processed using bcftools [44] version 1.6 to keep only called structural variants that passed Delly’s quality control.

Structural variants were then merged and filtered using SURVIVOR [45] version 1.0.7. Structural variants that were the same type, were on the same strand, and had breakpoints within 1000bp were merged. Only structural variants that were called by three or more callers and were larger than 50 bp were kept. Finally, the structural variants called for each dataset were all merged as before except breakpoints within 100bp were merged together. From this list of all structural variants, only genomic deletions were kept for further analysis.

### Definition of PAV orthogroups and conserved groups

For each lineage, PAV orthogroups were defined by first taking the matrix of validated PAVs and filtering this matrix to orthogroups that were present in at least two isolates and absent in at least two isolates. The SCO phylogeny of the lineage was then analyzed for each candidate PAV orthogroup. If the orthogroup was only absent in strains that formed a monophyletic group, the orthogroup was not considered to be a PAV orthogroup. Additionally, if the orthogroup was only found in strains that formed a monophyletic group, the orthogroup was not considered to be a PAV orthogroup either. All orthogroups that were therefore present in two independent groups and absent in two independent groups were labeled PAV orthogroups. All orthogroups that were missing one or fewer strains were considered conserved orthogroups. All other orthogroups were considered “other”.

### Transposable element annotation

TE annotation was performed using RepeatMasker [46] version 4.1.1 and a reference TE library for all pathotypes of *M. oryzae* generated by Nakamoto *et al*. [20]. The parameters -cutoff 250, -nolow, -no_is, and -norna were used for the RepeatMasker command.

### Next-generation sequencing data and GC content analysis

RNAseq data for MoO was obtained from SRA (Additional File 13) from a previous study [47] and mapped to the *M. oryzae* Guy11 genome (GCA002368485.1) for in culture data and the *M. oryzae* Guy11 genome combined with the *Oryza sativa* Nipponbare genome (GCA001433935.1) for the *in planta* data. RNAseq data for MoT was obtained from SRA accessions SRR9127598 through SRR9127602 from a previously published study [48] and mapped to the *M. oryzae* B71 genome (GCA004785725.2) for in culture data and the *M. oryzae* B71 genome combined with the wheat *Triticum aestivum* genome (GCA900519105.1) for the in planta data. Mapping was performed using STAR [49] version 2.7.1a and index files for mapping were made using the previously mentioned genomes and genome combinations along with corresponding gene annotation files obtained from FunGAP for the *M. oryzae* genomes, or from GenBank for the rice and wheat genomes. Read counts for each gene were calculated using the – quantMode GeneCounts parameter in STAR. These read counts were normalized to gene size as reads per kilobase values (RPK), then the total number of RPKs were summed for each sample and divided by one million. This sum was used to normalize read counts in each sample to obtain transcript per million (TPM) values for each sample. These TPM values were then averaged across treatments.

Published ChIPSeq data for H3K27me3, H3K27ac and H3K36me3 histone marks were obtained from a study published by Zhang *et al*. [47]. Published eccDNA sequencing data were obtained from a previous study by Joubert and Krasileva [50]. Reads were mapped to the *M. oryzae* Guy11 genome using BWA MEM [40] version 0.7.17-r1188. Read counts per gene were obtained using the coverage command from the BEDtools suite of tools [51] version 2.28.0. Read counts were normalized for gene and library size and averaged per treatment as for RNAseq data.

Methylation data from *M. oryzae* mycelium was obtained from a previous study published by Jeon *et al*. [52]. Reads were mapped to the *M. oryzae* genome and processed using the Bismark pipeline [53] version 0.24.0. Methylation percentage for all cytosines were extracted while ignoring the first 2 bases of all reads. The percentage of methylated cytosines was then calculated for a gene by averaging the methylation percentage of all cytosines in that gene.

To assign signal from next-generation sequencing datasets to orthogroups, signal for all orthologs in *M. oryzae* Guy11 and *M. oryzae* B71 within each orthogroup were averaged. Any orthogroups that did not have orthologs from B71 and Guy11 within them were given a value equal to the median value for all other orthogroups. *M. oryzae* Guy11 was not included in the original orthogrouping so a separate set of orthogroups were generated which included the *M. oryzae* Guy11 proteome annotated using FunGAP [24] as previously described in order to transfer next-generation sequencing signals.

Finally, GC content values for genes and flanking regions were calculated using the nuc command in BEDTools [51] version 2.28.0.

The same methods were used for calculating these values for genomic deletions.

### Profile plots

10bp windows were first generated for each *M. oryzae* reference genome. The number of TEs and the number of genes in each window were then calculated using the coverage command in BEDTools [51] version 2.28.0 and stored as bedgraph files. Bigwig files were generated from bedgraph files using the bedGraphToBigWig tool (https://www.encodeproject.org/software/bedgraphtobigwig/) version 4. Finally, data for profile plots of genomic deletions were generated using the computeMatrix scale-regions and the plotProfile commands of the DeepTools suite of tools [54] version 3.5.1.

### Random forest classification and feature importances calculation

Random forest classifiers were trained and performance statistics were calculated using the scikit-learn Python module [55] version 1.1.1. The hyperparameters used to train the model were as follows: 2000 estimators, a minimum of two samples to split a node, no minimum number of samples per leaf, no maximum tree depth, no maximum number of features per tree, and bootstrapping enabled. Classifiers were trained only on data for genes belonging to lineages 2 and 3 for MoO. Before training, all genes belonging to four genomes from each lineage were removed. From the remaining data, 50% of the genes not labeled as PAV were removed to improve the balance between PAV genes and non-PAV genes in the training data. The model was then trained and tested on the genes from the eight genomes that were removed before testing. The training and testing data split was repeated 100 times to generate average precision, recall, and F1 values as well as average number of true positives, false positives, true negatives, and false negatives for all models.

Feature importances were calculated according to methods described within the rfpimp Python module (https://github.com/parrt/random-forest-importances). Briefly, a random forest classifier was trained and tested as before to measure a baseline F1 statistic. Each variable in the testing data was then permuted in turn and a new F1 statistic for the model was generated on the permuted data. The difference between the baseline F1 and the new F1 were then calculated. This process was then repeated 100 times and the average decrease in the F1 statistic when each variable was permuted were reported.

Spearman and point biserial correlation coefficients between variables were calculated using the cor function in R version 3.6.1. Phi correlation coefficients were calculated using the psych package [56] version 2.2.9. To calculate dependence statistics for each variable in the complete MoO model, a random forest classifier or a random forest regressor was used to predict each variable originally used to train the PAV gene prediction model using all remaining variables. The same hyperparameters and train-test split were used to train and test each model as for the original PAV gene prediction model. Baseline F1 or R^2^ values for each model were then calculated and the change in these values when each variable within the model was permuted were calculated as before. However, the results reported were only from a single run of this analysis.

## Data processing and analysis

Data processing was performed in a RedHat Enterprise Linux environment with GNU bash version 4.2.46(20)-release. GNU coreutils version 8.22, GNU grep version 2.20, GNU sed version 4.2.2, gzip version 1.5, and GNU awk version 4.0.2 were all used for processing and handling. Conda (https://docs.conda.io/en/latest/) was used to facilitate installation of software and packages. Code parallelization was performed with GNU parallel [57] version 20180322. Previously published data was downloaded using curl version 7.65.3 (https://curl.se/) and sra-tools version 2.10.4 (https://github.com/ncbi/sra-tools). BED format files were processed using BEDtools [51] version 2.28.0. VCF format files were processed using bcftools [44] version 1.6. SAM and BAM format files were processed using SAMtools [44] version 1.8. FASTA format files were processed using seqtk (https://github.com/lh3/seqtk) version 1.2-r102-dirty.

Data processing and analysis were performed using custom Python scripts written in Python version 3.10.5 with the help of pandas [58] version 1.4.3 and numpy [59] version 1.23.1. GFF format files were parsed in Python using BCBio GFF version 0.6.9 (https://github.com/chapmanb/bcbb/tree/master/gff). FASTA format files were processed in python using SeqIO from Biopython [60] version 1.80.

Data processing and analysis were also performed using custom R scripts written in R version 3.6.1 with the help of data.table [61] version 1.13.6, tidyr [62] version 1.1.3, reshape2 [63] version 1.4.4, and dplyr [64] version 1.0.4. Plotting was performed using the ggplot2 package [65] version 3.3.5 and the ggnewscale package [66] version 0.4.8. Phylogenies were analyzed and plotted using the ape [31] package version 5.5 and the phytools package [67] version 0.7.90.

## Declarations

### Availability of data and materials

All code for data generation, analysis, and plotting is available on GitHub: https://github.com/pierrj/moryzae_pav_manuscript_code

All files used for analysis and plotting are available on Zenodo under the DOI 10.5281/zenodo.7444379.

### Consent for publication

Not applicable.

### Ethics approval and consent to participate

Not applicable.

### Authors’ contributions

PMJ conceptualized and designed the study with input from KVK. PMJ designed the analyses and analyzed the data. PMJ wrote the original draft manuscript. PMJ and KVK reviewed and edited the manuscript. Both authors read and approved the final manuscript.

## Acknowledgements

We thank Krasileva lab members for thoughtful comments and feedback on the manuscript. This research used the Savio computational cluster resource provided by the Berkeley Research Computing program at the University of California, Berkeley (supported by the UC Berkeley Chancellor, Vice Chancellor for Research, and Chief Information Officer).

## Supplementary Information

Additional File 1: Supplementary Figures.

Fig. S1. Phylogeny of rice-infecting *M. oryzae* isolates used in this study.

Fig. S2. Phylogeny of wheat-infecting *M. oryzae* isolates used in this study.

Fig. S3. Distances to the nearest gene for PAV and conserved genes in MoO and MoT. A. Density plots showing the distribution of the distances to the nearest gene for conserved and PAV genes in MoO and MoT.

Fig. S4. Profile plots showing transposable element (TE) and gene density within genomic regions of the rice and wheat-infecting *M. oryzae* genomes.

Fig. S5. Density plots of additional features of PAV and conserved genes.

Fig. S6. Comparison of various functional annotations of PAV and conserved genes.

Fig. S7. Density plots showing the distributions of various features of MoO and MoT genomic deletions.

Fig. S8. Correlation coefficients for variables included in the MoO random forest classifier.

Fig. S9. Dependence matrix of variables included in the MoO random forest classifier.

Fig. S10. Confusion matrices for the MoT random forest classifier and the MoO random forest classifier trained on a subset of features.

Additional File 2: List of accessions for MoT genomes.

Additional File 3: List of accessions for MoO Illumina data.

Additional File 4: List of accessions for MoT Illumina data.

Additional File 5: List of genomic deletions called using MoO Illumina sequencing data.

Additional File 6: List of genomic deletions called using MoT Illumina sequencing data.

Additional File 7: Table of median upstream and downstream distances to nearest PAV gene, TE, and gene for MoO and MoT. The p-values shown are two-tailed p-values resulting from permutation tests for the differences in medians between PAV and conserved genes for each pathotype with 1,000 permutations.

Additional File 8: Table showing the number of PAV and conserved genes that are near PAV genes, near TEs, near genes, have a TM domain, have a signal peptide, are predicted effectors, have a GO annotation, and have a PFAM domain annotation for MoO and MoT. The p-values shown were the results of Chi-squared tests used to test for indepedence between the PAV/conserved gene label and each feature for each pathotype.

Additional File 9: Table showing the number of PAV and conserved genes that are near PAV genes, near TEs, near genes, have a TM domain, have a signal peptide, are predicted effectors, have a GO annotation, and have a PFAM domain annotation for MoO and MoT. The p-values shown were the results of Chi-squared tests used to test for indepedence between the pathotype and each feature for each PAV/conserved gene label.

Additional File 10: Table showing the mean, median, standard deviation, 25^th^ percentile and 75^th^ percentile for the distributions of various continuous variables that describe PAV and conserved genes in MoO and MoT. The p-values shown are two-tailed p-values resulting from permutation tests for the differences in each statistic between PAV and conserved genes for each pathotype with 1,000 permutations.

Additional File 11: Table showing the mean, median, standard deviation, 25^th^ percentile and 75^th^ percentile for the distributions of various continuous variables that describe PAV and conserved genes in MoO and MoT. The p-values shown are two-tailed p-values resulting from permutation tests for the differences in each statistic between pathotypes for PAV and conserved genes with 1,000 permutations.

Additional File 12: Table showing the mean, median, standard deviation, 25^th^ percentile and 75^th^ percentile for the distributions of various continuous variables that describe genomic deletions and baseline genomic regions in MoO and MoT. The p-values shown are two-tailed p-values resulting from permutation tests for the differences in each statistic between deletions and baseline for each pathotype with 1,000 permutations.

Additional File 13: List of accessions for MoO RNAseq data.

## Additional Files

Additional File 1: Supplementary Figures.

**Fig. S1.**
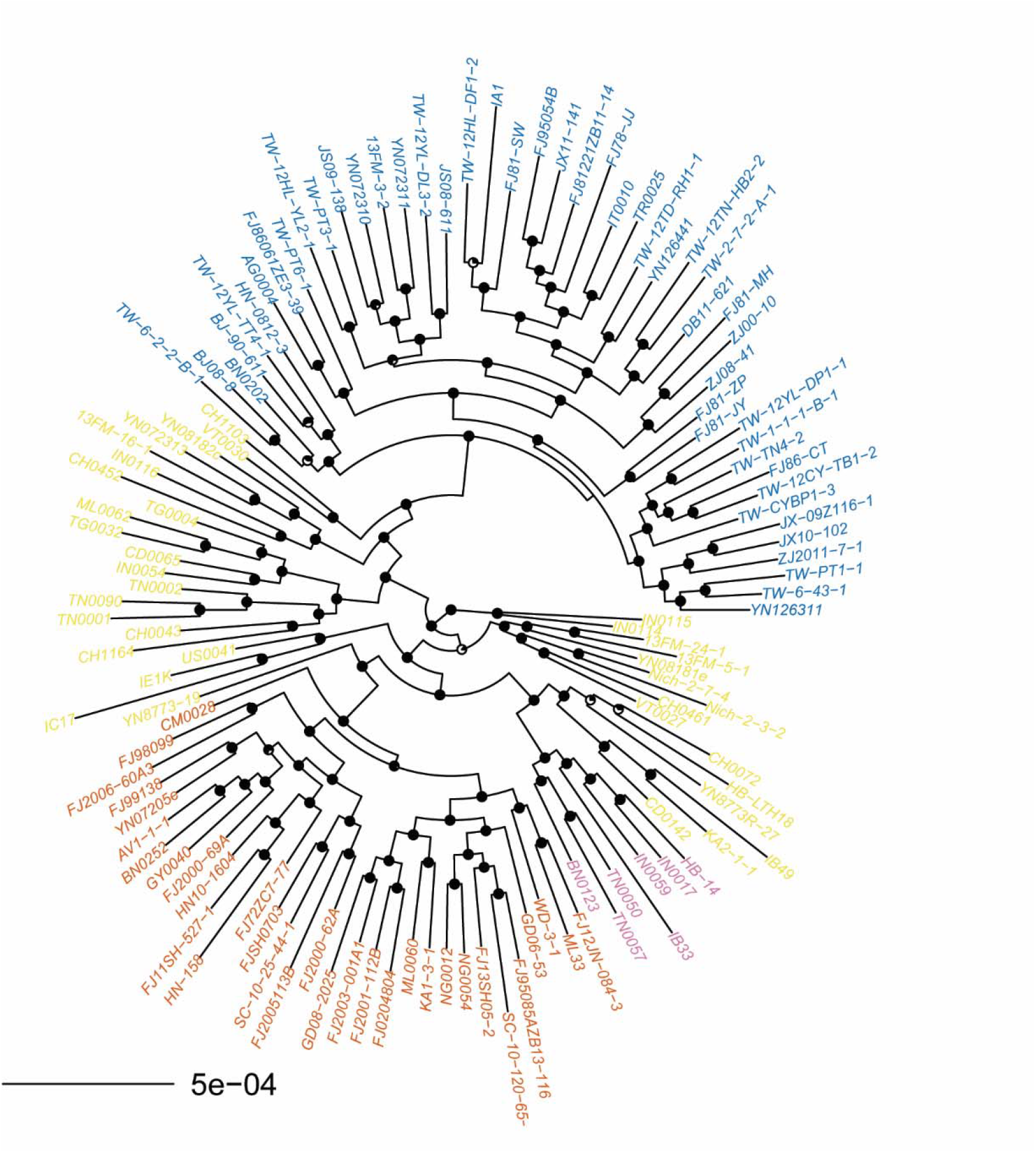
Phylogeny of rice-infecting *M. oryzae* isolates used in this study. Phylogeny was generated using a multiple-sequence alignment of SCOs and fasttree [30]. Pie charts on nodes represent the fraction of bootstrap replicates that support the node. Isolates belonging to lineage 1 are colored yellow, isolates belonging to lineage 2 are colored orange, isolates belonging to lineage 3 are colored blue, and isolates belonging to lineage 4 are colored pink. Lineages were named as previously described [17].

**Fig. S2.**
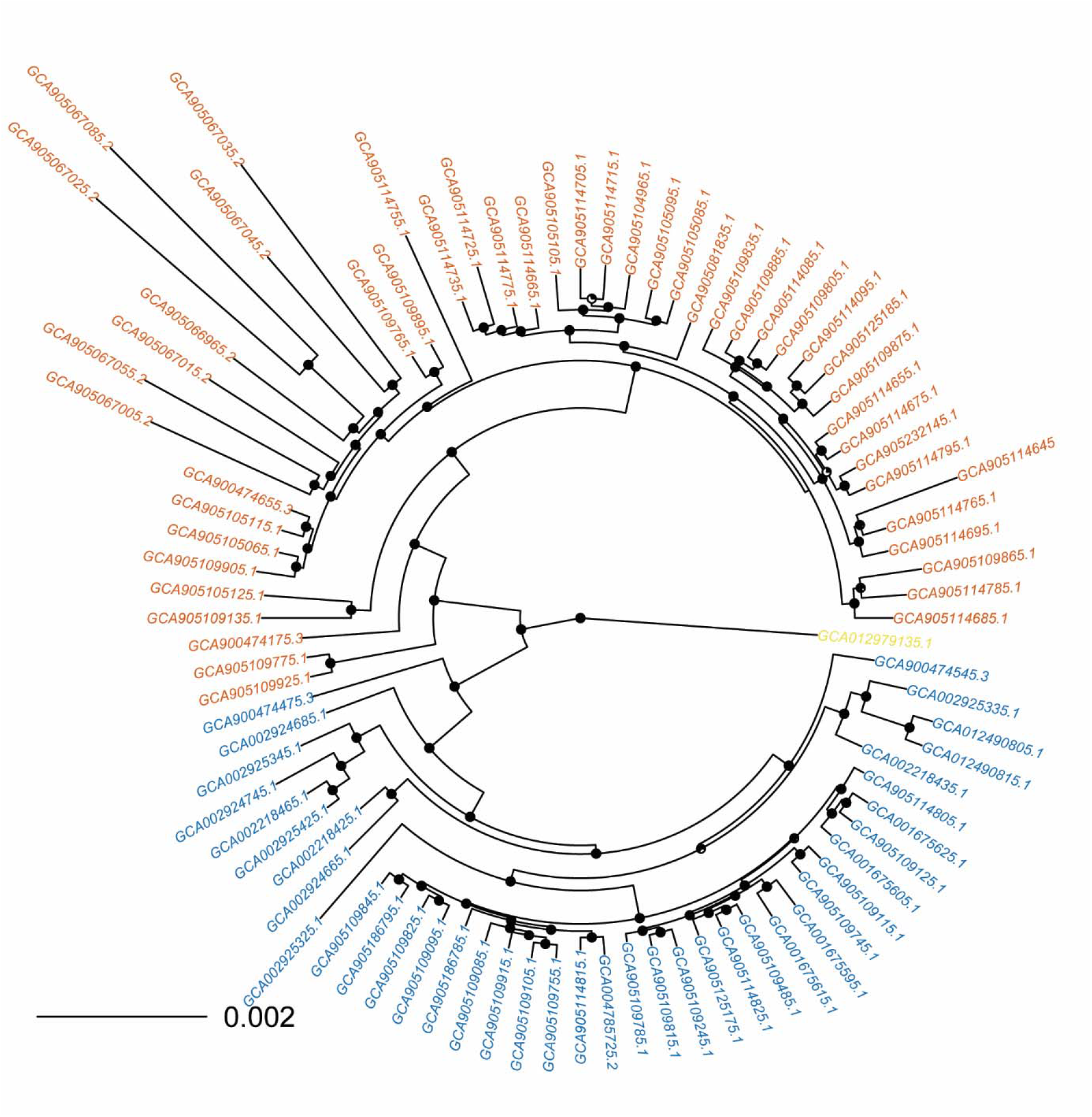
Phylogeny of wheat-infecting *M. oryzae* isolates used in this study. Phylogeny was generated using a multiple-sequence alignment of SCOs and fasttree [30]. Pie charts on nodes represent the fraction of bootstrap replicates that support the node. Isolates belonging to lineage 1 are colored orange, isolates belonging to lineage 2 are colored blue, and the isolate that was not assigned to either lineage is colored yellow.

**Fig. S3.**
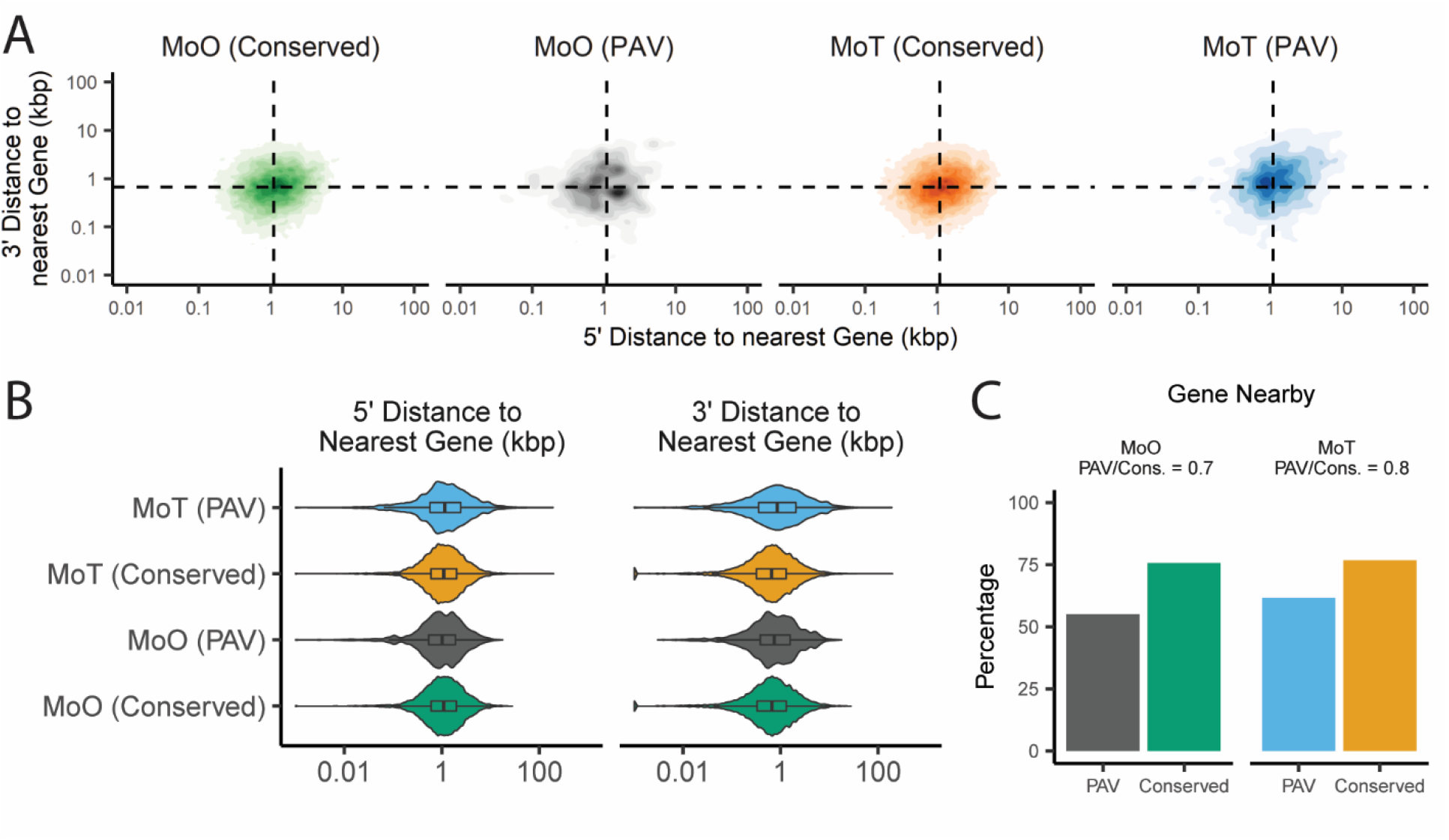
Distances to the nearest gene for PAV and conserved genes in MoO and MoT. A. Density plots showing the distribution of the distances to the nearest gene for conserved and PAV genes in MoO and MoT. B. Violin plot showing the distribution of the distances to the nearest gene for conserved and PAV genes in MoO and MoT. C. Percentages and proportions of PAV and conserved genes that are within 1000bp of another gene in MoO and MoT. Dashed lines in density plots represent the median values for all genes in both pathotypes. Median values and statistical comparisons for data shown are listed in Additional File 7, Additional File 8, and Additional File 9.

**Fig. S4.**
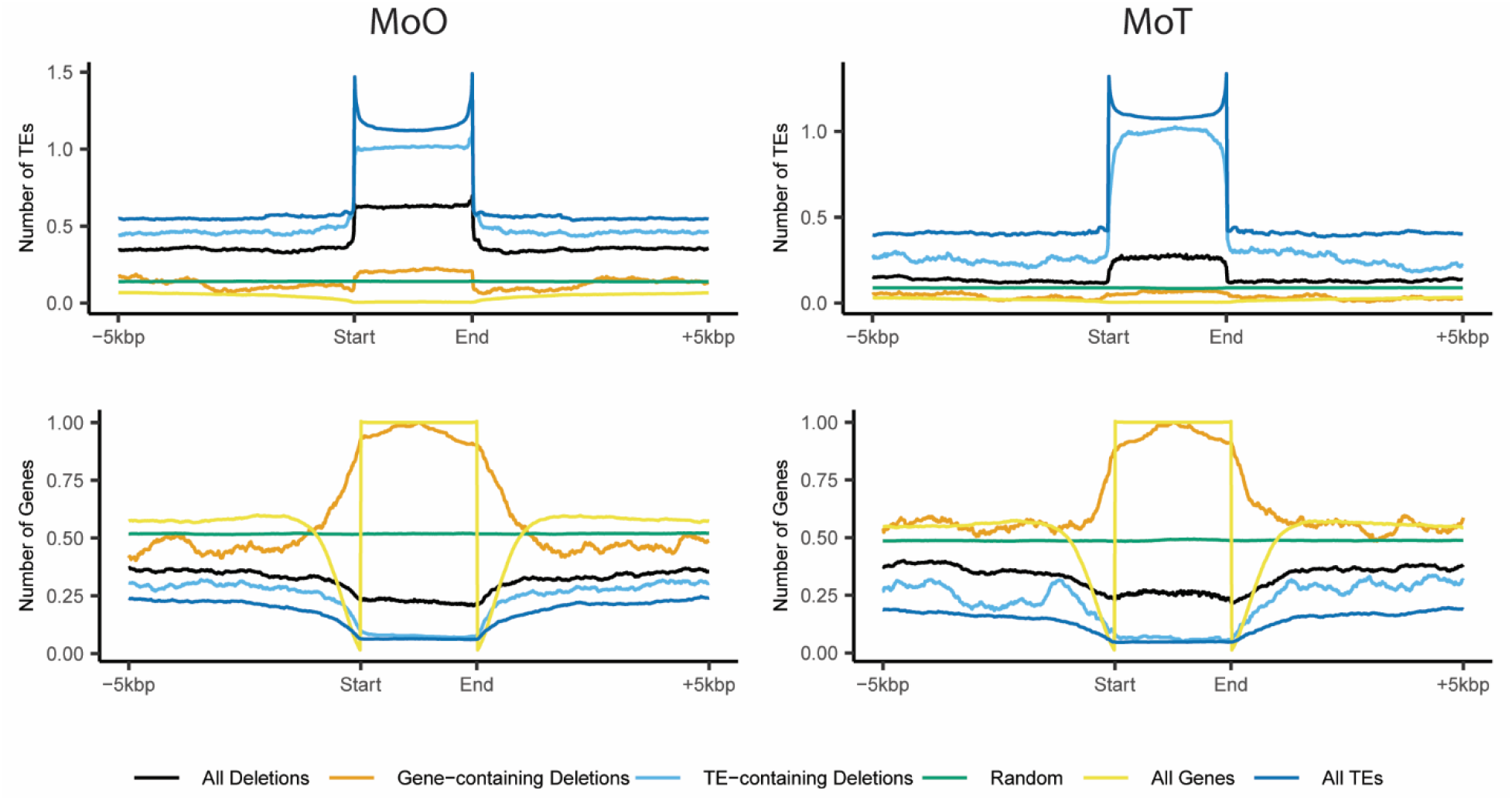
Profile plots showing transposable element (TE) and gene density within genomic regions of the rice and wheat-infecting *M. oryzae* genomes. The flanking regions of these regions are also shown. Gene- and TE-containing regions represent the subet of all deletions that overlapped at least 50% with a gene or TE sequence, respectively. Genomic deletions were shuffled throughout the genome 100 times to generate the data for random regions in the plots.

**Fig. S5.**
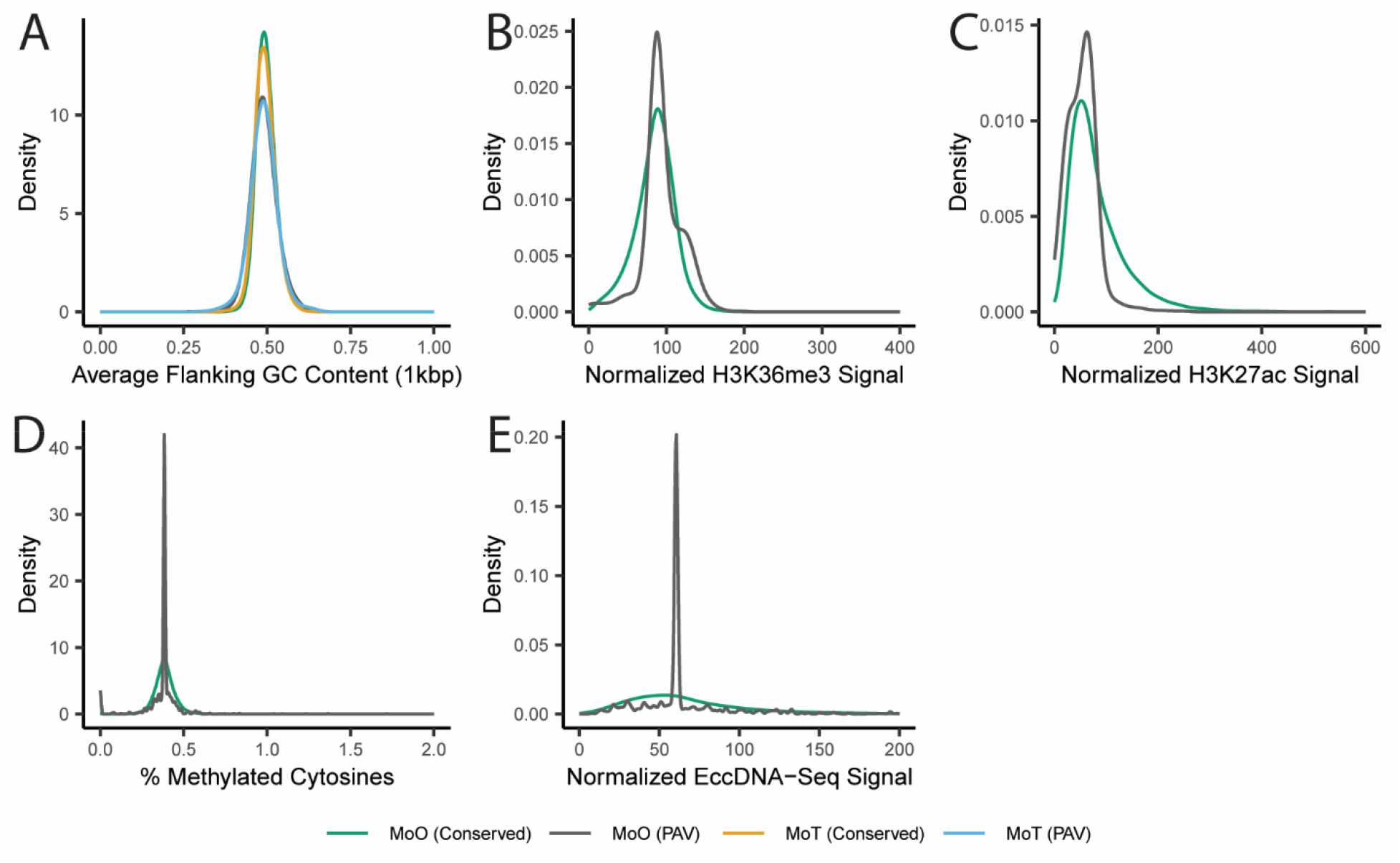
Density plots of additional features of PAV and conserved genes. Density plots showing the distributions of A. average flanking GC content, B. normalized H3K36me3 histone mark ChIP-Seq signal, C. noramlized H3K27ac histone mark ChIP-Seq signal, D. average % methylation of cytosines, and E. normalized extrachromosomal DNA (eccDNA) sequencing signal for PAV and conserved genes in MoO and MoT. In panel A, the line representing the data for MoO PAV genes appears behind the line representing data for MoT PAV genes. In panels B, C, D, and E, MoT genes were not included as this data is not available for MoT. Statistics describing distributions and statistical comparisons between these statistics are listed in Additional File 10 and Additional File 11.

**Fig. S6.**
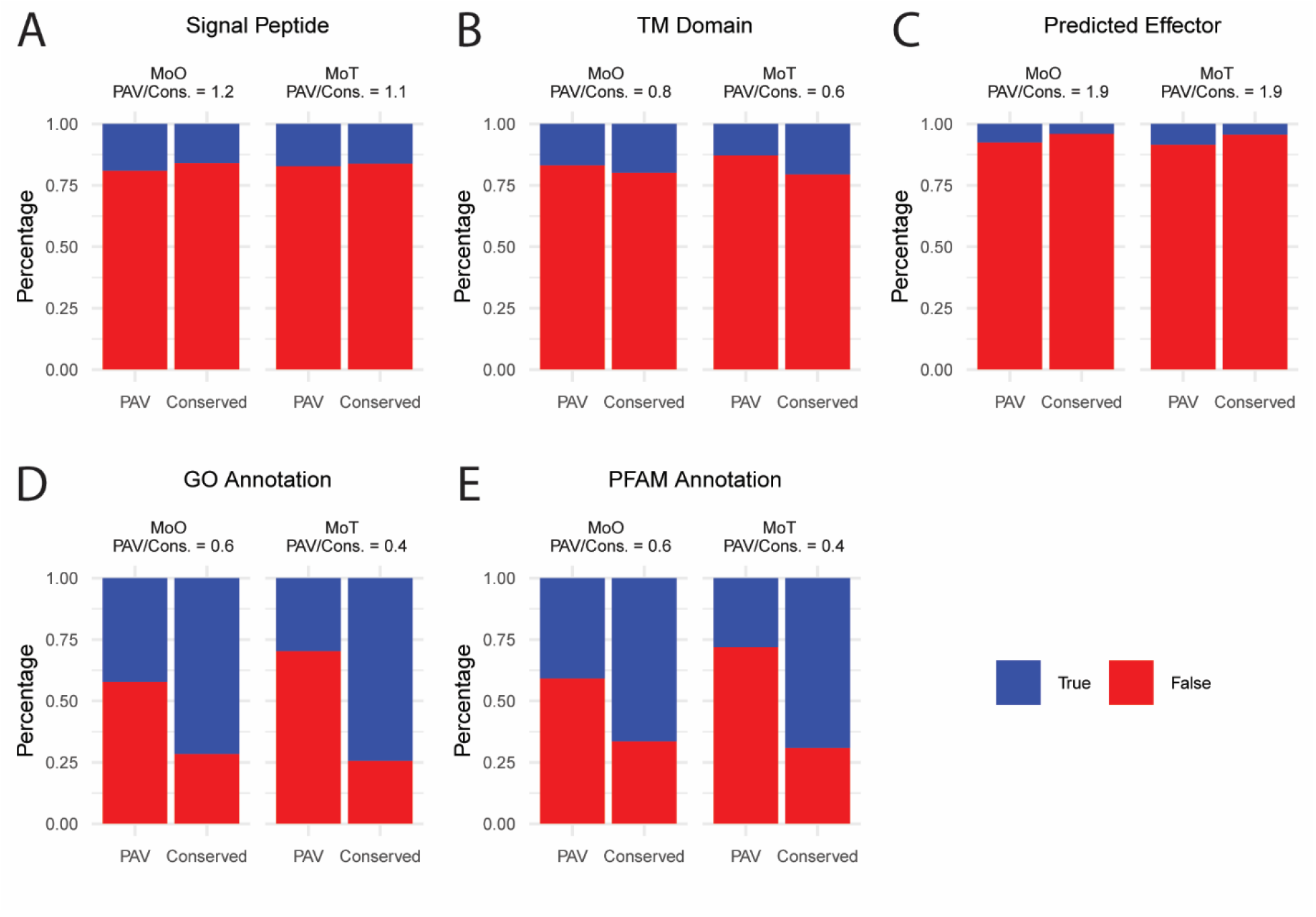
Comparison of various functional annotations of PAV and conserved genes. Comparison of percentages and ratios of PAV and conserved genes annotated as A. having a signal peptide, B. having a transmembrane (TM) domain, C. being a predicted effector, D. having a GO annotation, and E. having a protein family (PFAM) domain annotation for MoO and MoT genes. Counts for each category and stastical comparisons of these counts are listed in Additional File 8 and Additional File 9.

**Fig. S7.**
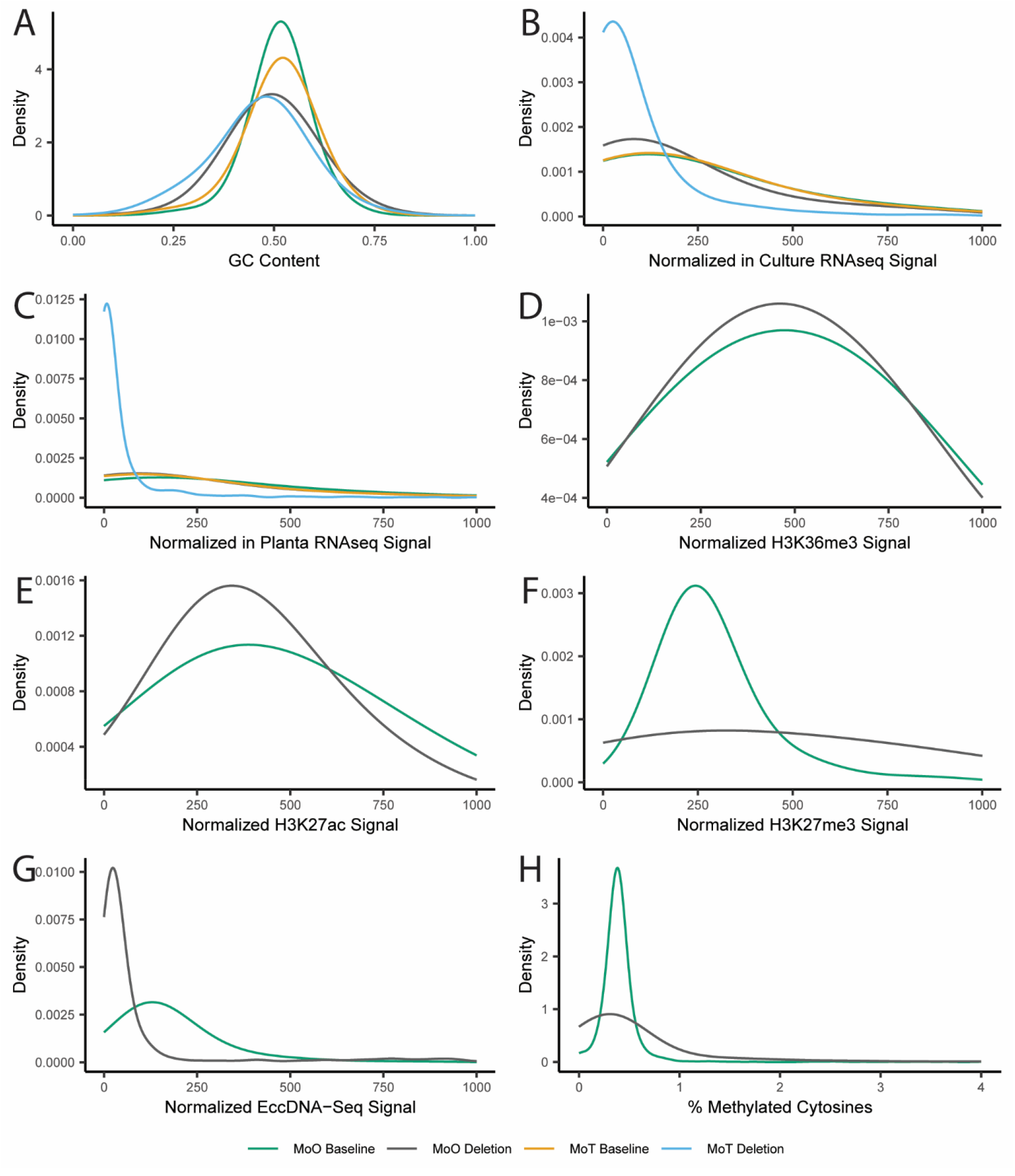
Density plots showing the distributions of various features of MoO and MoT genomic deletions. Density plots showing the distributions of A. AT content, B. normalized in culture RNAseq signal, C. normalized in planta RNAseq signal, D. normalized H3K36me3 histone mark ChIP-Seq signal, E. normalized H3K27ac histone mark ChIP-Seq signal, F. normalized H3K27ac histone mark ChIP-Seq signal, G. normalized eccDNA sequencing signal, and H. average % methylation of cytosines for genomic deletions in MoO and MoT, as compared to baseline. Genomic baseline values were generated by shuffling the deletions throughout the portions of the genome that were not deleted in any isolate. Statistics describing distributions and statistical comparisons between these statistics are listed in Additional File 12.

**Fig. S8.**
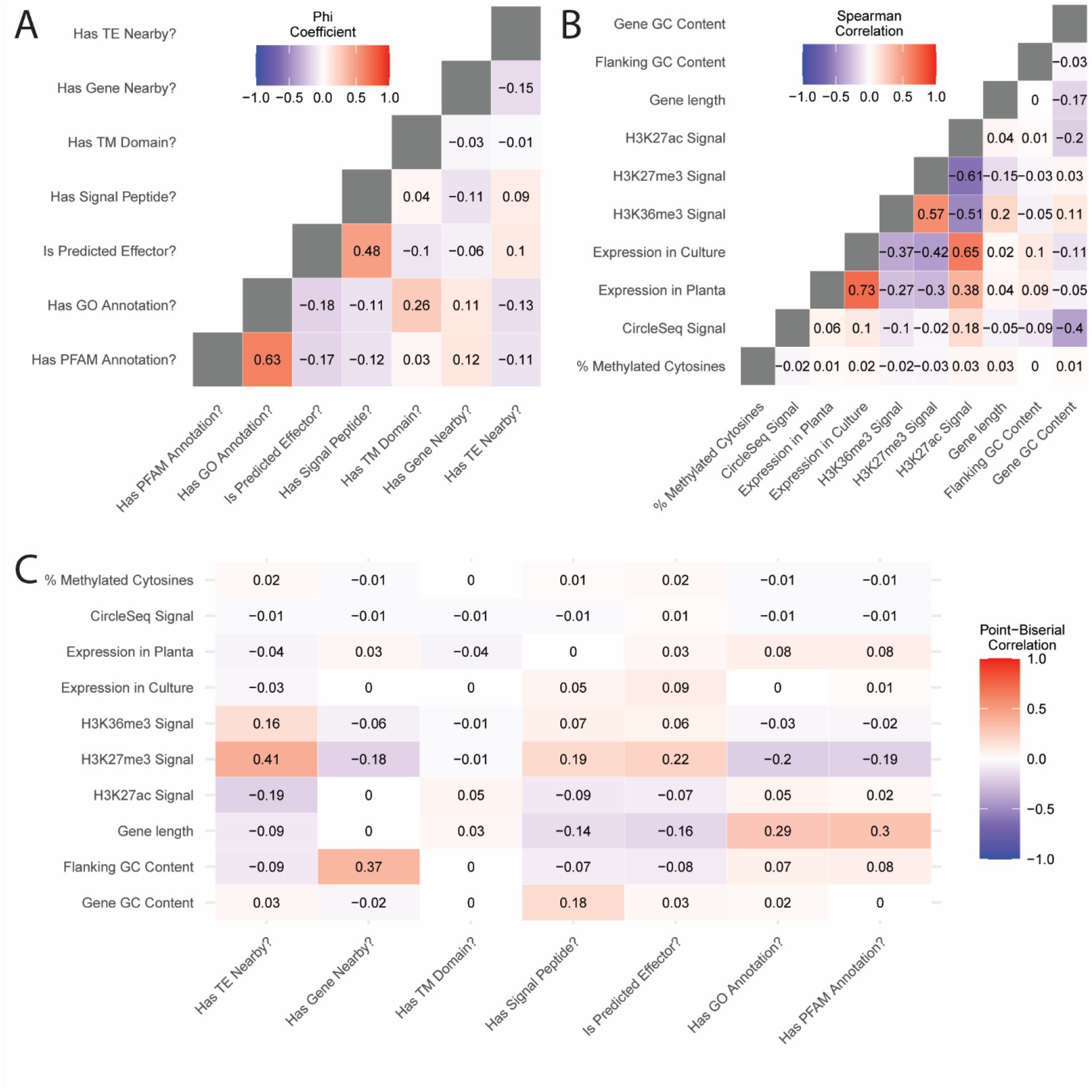
Correlation coefficients for variables included in the MoO random forest classifier. Heat map representing A. Phi coefficient between binary variables, B. Spearman rank correlation coefficient between continuous variables, and C. Point-Biserial correlation coefficient between continuous and binary variables.

**Fig. S9.**
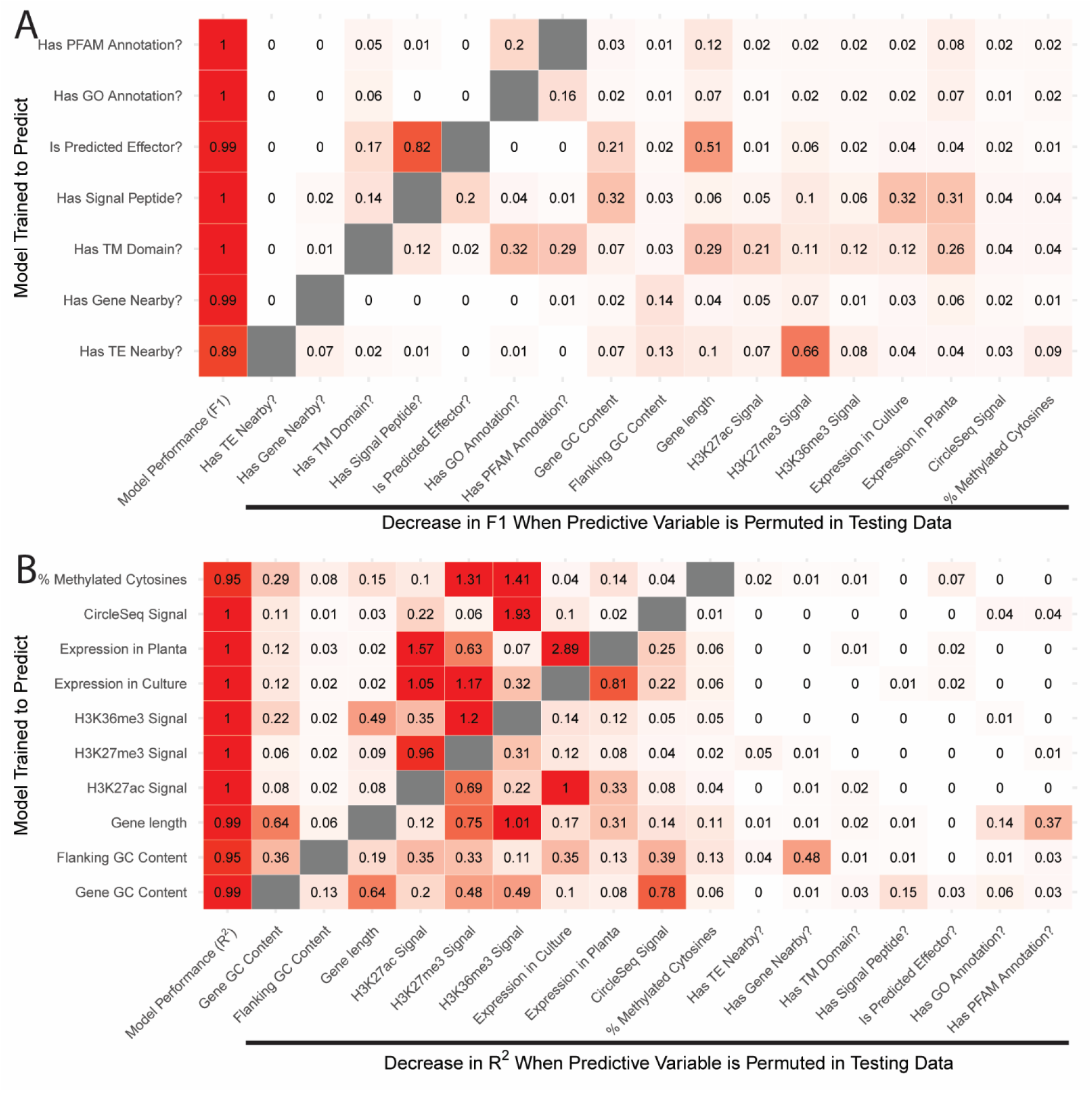
Dependence matrix of variables included in the MoO random forest classifier. A model was trained to predict each variable used in our MoO random forest classifier using the remaining variables. A. Heatmap representing the F1 statistic of each model when trained to predict categorical variables and decrease in F1 when predictive variables were permuted in the testing data. B. Heatmap representing the R^2^ statistic of each model when trained to predict categorical variables and decrease in R^2^ when predictive variables were permuted in the testing data.

**Fig. S10.**
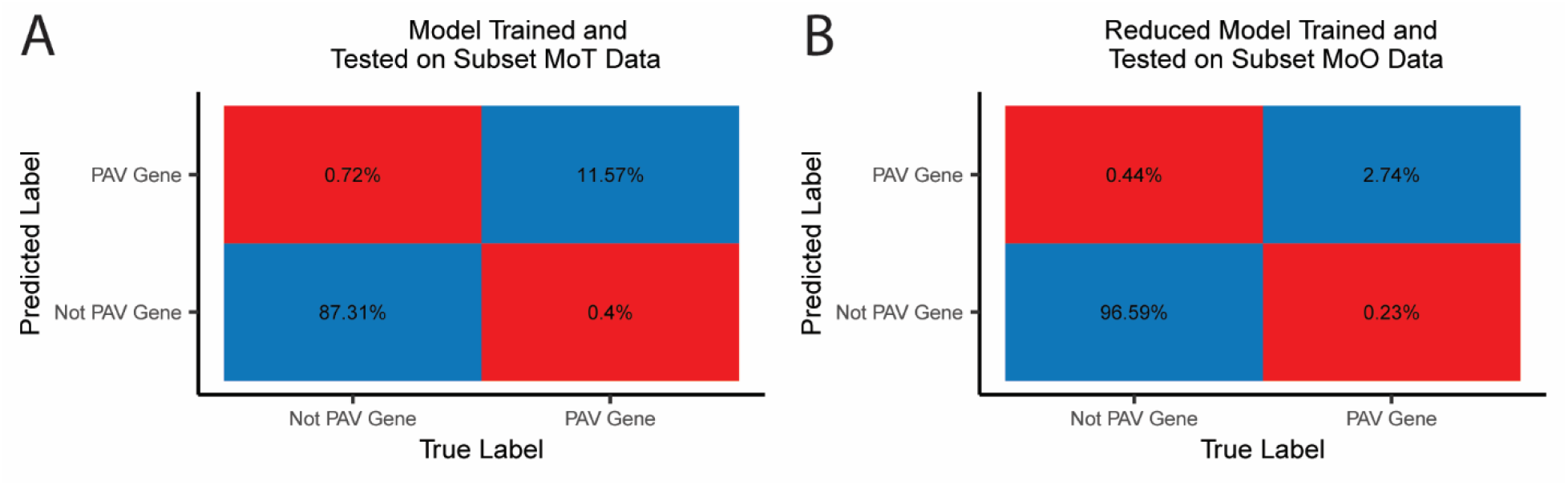
Confusion matrices for the MoT random forest classifier and the MoO random forest classifier trained on a subset of features. A. Confusion matrix showing predictions of the MoT random forest classifier when tested on MoT genes that it was not trained on. B. Confusion matrix showing predictions of the MoO random forest classifier trained on a subset of features when tested on MoO genes that it was not trained on.

